# Rerouting of ribosomal proteins into splicing in plant organelles

**DOI:** 10.1101/2020.03.03.974766

**Authors:** Chuande Wang, Rachel Fourdin, Martine Quadrado, Céline Dargel-Graffin, Dimitri Tolleter, David Macherel, Hakim Mireau

## Abstract

Production and expression of RNAs requires the action of multiple RNA-binding proteins (RBPs). New RBPs are most often created by novel combinations of dedicated RNA binding modules. However, recruiting existing genes to create new RBPs is also an important evolutionary strategy. In this report, we analysed the 8-member uL18 ribosomal protein family in Arabidopsis. uL18 proteins share a short structurally conserved domain that binds the 5S rRNA and allow its incorporation into ribosomes. Our results indicate that Arabidopsis uL18-like proteins are targeted to either mitochondria or chloroplasts. While two members of the family are found in organelle ribosomes, we reveal that two uL18-type proteins correspond to splicing factors that are necessary for the elimination of certain mitochondrial and plastid group II introns. These two proteins do not co-sediment with mitochondrial or plastid ribosomes but associate with the introns whose splicing they promote. Our study thus reveals that the RNA binding capacity of uL18 ribosomal proteins has been detoured to create factors facilitating the elimination of organellar introns.

## Introduction

The evolution of eukaryotic cells has involved the acquisition of highly specialized energy-producing organelles such as mitochondria and plastids, both of which originate from the adoption of bacteria (1, 2). The transformation of the original endosymbionts into specialized organelles followed a similar trend comprising a massive reduction of their genomes and a progressive loss of autonomy. A large number of host-derived factors, either acquired or recruited during evolution, became essential for proper expression of the essential genes that are still present in modern mitochondrial and plastid genomes. The interplay between ancient bacterial-like infrastructures and eukaryotic-derived functions resulted in organellar gene expression mechanisms that are far more complex than that of modern bacteria (3, 4). An astonishing complexity has been reached for all processes dealing with the production and the expression of mitochondrial and plastid transcripts, especially in terrestrial plants. Higher plant mitochondrial and plastid genomes produce a complex array of mono and polycistronic transcripts that need extensive processing to accumulate. They also undergo extensive sequence modification through C-to-U RNA editing and contain multiple group II introns that need to be removed prior to translation. Each of these RNA processing steps requires the action of dedicated ribonucleoprotein complexes that are still poorly characterized. Several classes of nuclear-encoded RNA-binding proteins were found to play roles in mitochondrial and/or plastid RNA expression like the pentatricopeptide repeat (PPR) proteins (5, 6) and other kinds of helical repeat protein families (7) which all adopt similar solenoid like structures exposing key amino acids for RNA binding (7–9). Other protein families like RNA recognition motif (RRM), multiple organellar RNA editing factor (MORF), chloroplast RNA splicing and ribosome maturation (CRM), plant organellar RNA recognition (PORR) and maturase proteins play also role in plant organellar RNA expression (10–14) (15). Some of these factors were created using generic RNA binding domains (such as the RRM domain) while others evolved by hijacking proteins with other functions such as CRM proteins that are likely derived from a bacterial protein involved in ribosome maturation (11). Ribosomal proteins (RPs) represent another large class of conserved and abundant RNA binding proteins. Although their primary function is to stabilize ribosomal RNA structures to guarantee the efficiency of protein synthesis, extra-ribosomal employments have been described in a few cases (16–18). Plant ribosomal proteins are often encoded by small multigene families comprising up to eight active genes, suggesting potential specialization or extra-ribosomal activities for some family members (19, 20). In this study, we investigated the function of a small protein family in Arabidopsis whose members were annotated to be homologs of the uL18 ribosomal protein. The detailed analysis of two of these uL18 homologs revealed that they lost the ability to associate with ribosomes and have become organellar group II intron splicing factors.

## Materials and Methods

### Plant material

Arabidopsis (*Arabidopsis thaliana*) Col-0 plants were obtained from the Arabidopsis stock centre of the Institut National de Recherche pour l’Agriculture, l’Alimentation et l’Environnement in Versailles (http://dbsgap.versailles.inra.fr/portail/). The Arabidopsis *ul18-l1* (N581267) and *ul18-l8* (N568525 and CS841212) mutants were obtained from the SALK mutant collection (21). Plants were grown in a greenhouse in long day conditions for 10 to 12 weeks before use. Flowers or leaves were harvested simultaneously for all genotypes, snap-frozen in liquid nitrogen and stored at −80°C until use.

For the complementation assays, the coding sequence of *uL18-L1* or *uL18-L8* without stop codon but with 1 kb of promoter region was amplified by PCR and cloned to pGWB13 (22) by Gateway™ cloning, creating C-terminal translational fusions with the 3HA tag. The resulting plasmids were used to transform *ul18-l1* or *ul18-l*8-1 heterozygous plants by floral dipping and complemented homozygous mutant plants were identified in the progenies of transgenic plants.

### Preparation of organelles and stromal protein

Arabidopsis mitochondria and chloroplasts were purified from flower buds and young leaves, respectively, as described in (23). Stromal protein fractions were prepared from purified plastids as previously described in (24).

### Immunodetection of proteins

Proteins were extracted from flowers, leaves or purified organelles in 30 mM HEPES-KOH (pH 7.7), 10 mM Mg(OAc)_2_, 150 mM KOAc, 10% glycerol and 0.5% (w/v) CHAPS. Protein concentrations were measured with Bradford (Bio-Rad) and separated by SDS-PAGE. After electrophoresis, gels were transferred onto PVDF membranes (Perkin-Elmer) under semidry conditions. Membranes were hybridized with antibodies using dilutions indicated in Supplementary Table S2.

### Subcellular distributions of uL18-1 and uL18-8 proteins

DNA regions corresponding to the coding sequence of each uL18-Like protein but lacking the stop codon were PCR amplified and transferred into pGWB5 (22) by Gateway™ cloning (Invitrogen) to create GFP translational fusions. The constructs were used to transform Arabidopsis plants by floral dipping and GFP fluorescence was visualized in leaf or root cells by confocal microscopy. When necessary, roots of transgenic plants were soaked in a solution of 0.1 μM MitoTracker™ Red to label mitochondria prior to observation.

### RNA analysis

RNA extraction, northern blot and quantitative RT-PCR analyses were done as previously described in (25) with primer sets detailed in (26, 27).

### RNA immunoprecipitation assays

Immunoprecipitation of uL18L1-3HA and uL18L8-3HA were performed using the μMACS HA-Tagged Protein Isolation Kit (Miltenyi Biotec). Briefly, total proteins were extracted from 1 g of Arabidopsis transgenic cells expressing either uL18L1-3HA or uL18L8-3HA in 1 mL of cold lysis buffer (20 mM HEPES-KOH, pH 7.6, 100 mM KCl, 20 mM MgCl_2_, 1 mM DTT, 1% Triton X-100, 1X of complete EDTA-free protease inhibitor (Roche)) for 30 mins on ice. The lysates were clarified by centrifugation at 100,000 *g* for 20 mins at 4°C and resulting supernatants were incubated with 50 μL of anti-HA magnetic beads or protein A magnetic beads (Thermo Fisher Scientific) for 1 h at 4°C with rotation (10 rpm). After three washes with 1 mL of washing buffer (lysis buffer with only 0.1% Triton X-100), proteins were eluted in 120 μL of elution buffer provided by the kit and then subjected to immunoblot analysis. Co-immunoprecipitated RNAs were extracted with TRI Reagent**®** (Invitrogen) and treated with the RTS DNase I (Mobio) prior to RT-qPCR analysis. RNAs representing 1% of the input fraction and the totality of immunoprecipitated RNAs were used for cDNA synthesis. Three microliters of 10-fold diluted cDNA solutions were used in each qPCR reaction.

### Blue Native gel and in-gel activity assays

One hundred micrograms of mitochondrial proteins were separated on 4-16% (w/v) polyacrylamide Native PAGE gels (Invitrogen) and then either transferred to PVDF membranes or subjected to in-gel activity staining as in previously described in (25). Thylakoid membranes and BN-PAGE analysis were performed as described in (28), with minor modifications. After electrophoresis, the separated protein complexes were transferred to PVDF membranes for immunoblotting or stained in Coomassie blue R-250. Antibodies listed in Supplementary Table S2 were used for immuno-detection.

### Sucrose density gradient centrifugation

One gram of transformed Arabidopsis cells and the equivalent of 3 to 6 mg of stromal or total mitochondrial proteins were resuspended in 1 mL (for cells) or 200 μL (for organelles) of lysis buffer (30 mM HEPES-KOH (pH 7.8), 100 mM KOAc, 10 mM Mg(OAc)_2_, 5 mM β-mercaptoethanol, 5% glycerol, 1% n-dodecyl β-D-maltoside [DDM] and 1X cOmplete™ EDTA-free protease inhibitor (Roche)) and incubated at 4°C for 30 mins. The lysates were then diluted with the same volume of lysis buffer with no DDM and then cleared by ultracentrifugation at 100,000 *g* for 20 mins. Volumes corresponding to 2 mg of proteins were layered on continuous sucrose gradients (10%-30%) made in lysis buffer without DDM. Sucrose gradients were ultracentrifuged at 268,000 *g* for 4 hours at 4°C. Gradients were fractionated into 16 fractions of 330 μL and subsequently analysed by immunoblotting.

### Transformation of Arabidopsis cell suspension cultures

The coding sequences of *uL18-L1* and *uL18-L8* without their stop codon were PCR amplified and transferred by Gateway™ cloning (Invitrogen) into pGWB14 (22) resulting in C-terminal fusions with the 3xHA coding sequence, expressed under the control of the 35S promoter. These constructs were used to transform an Arabidopsis cell culture as indicated in (29).

### Analysis of photosynthesis and pigment composition

Plants were grown in long-day conditions (16:8 hours, 19:21°C L:D) at 120 μmol photons.m^−2^.s^−1^ for 4 weeks. Water content of rosette leaves was estimated by measuring the fresh and dry weight (after 48 h at 90°C) of detached rosettes. Leaf chlorophyll and carotenoid pigments were extracted with dimethylformamide and measured according to (30). Chlorophyll fluorescence of rosettes was measured non-invasively at room temperature with a PSI Open FluorCam FC 800-O (Photon Systems Intruments). For each plant, 13 spots, one per leaf (9 spots of 0.02 mm^2^ about 500 pixels and 4 spots of 0.01 mm^2^ about 200 pixels) were used to get a spatial average of fluorescence over time. Maximum fluorescence was obtained by saturation pulses at 2,800 μmol photons m^−2^.s^−1^ for 600 ms. Chlorophyll fluorescence parameters (F_v_/F_m_, 1-qP, qL, NPQmax, NPQrelax) were calculated from F_m’_ (F_m_ if after 45 mins of dark adaption), F_0_’ (F_0_ if after 45 mins of dark adaption) and F as maxima, minimal and current fluorescence respectively. Actinic light (420 μmol photons.m^−2^.s^−1^) was turned on for 500 s (with a saturating pulses every 80 s) and then dark relaxation time for 280 s (with saturating pulse at 80 s and then 200 s later). The maximum quantum efficiency of PSII photochemistry, Fv/Fm, was calculated as (Fm-Fo)/Fm, the Photochemical quenching, 1-qP as 1-(F_m’_-F)/(F_m’_-F_0’_), the fraction of open PSII centres, qL as (F_m’_-F)/(F_m’_-F_0’_), and the Non Photochemical Quenching, NPQ as (F_m_-F_m’_)/F_m’_ with NPQ_max_ just before turning the light off and NPQ_relax_ at the end of the 280 s of relaxation period of dark.

## Results

### The uL18 protein family comprises eight members in Arabidopsis

While analysing sequence variability among ribosomal proteins in Arabidopsis, we noticed that eight proteins were classified in the L18pL5e superfamily. These proteins share a rather degenerate but structurally conserved C-terminal RNA binding domain forming three α-helices and three and a half β-sheets that was shown to bind the 5S rRNA (31) (Figure 1). C-terminal GFP translational fusions with each of these uL18 like proteins revealed that five members are targeted to mitochondria and three to plastids (Figure 2). Multiple sequence alignments indicated that uL18-like members are more homologous to the *E. coli* uL18 protein than to the cytosolic uL18, strongly suggesting a bacterial origin for the uL18-like family (Supplementary Figure S1). Moreover, two members of the uL18 family, AT1G48350 and AT5G27820, have sequences and sizes very close to the *E. coli* uL18 protein. A protein 95% identical to AT1G48350 was found in the structure of the spinach (*Spinacia oleracea*) chloroplast ribosome (32), indicating that it most likely corresponds to the plastid uL18 protein in Arabidopsis. We thus named this protein uL18c in agreement with the new ribosomal protein nomenclature (33). Similarly, the AT5G27820 protein was recently found to be a mitoribosomal subunit (34) and was thus denominated uL18m (Figure 1). Our GFP fusion analysis confirmed the plastid and mitochondrial targeting of Arabidopsis uL18c and uL18m, respectively (Figure 2). The function of the six other proteins could not be recognized from their sequence and were named uL18-Like (or uL18-L) proteins. Potential orthologs could be easily identified for most uL18-Like members in both monocots and dicots, supporting their essential character for organelle functions (Supplementary Figure S1). The only exception is the uL18-L6 protein, which appears to be absent in monocotyledons.

**Figure 1:**
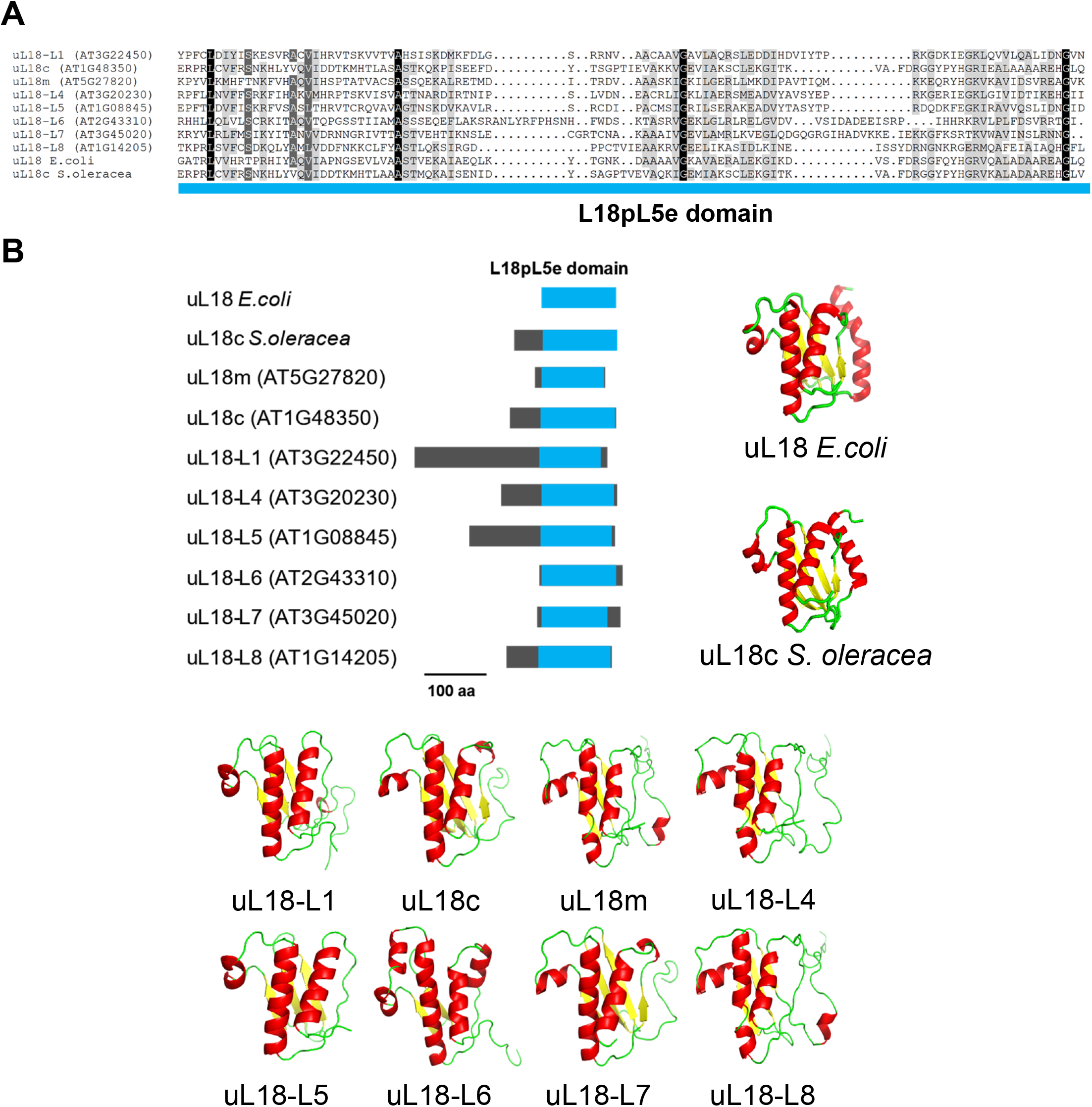
The Arabidopsis nuclear genome encodes 8 different uL18-like proteins. (A) Sequence alignment of the C-terminal L18pL5e domain of the eight uL18-like Arabidopsis proteins, the uL18 protein from *E. coli* and the uL18c from *S. oleracea.* The residues are colored according to the percentage of conservation from dark grey (100% identical) to light grey (50 % identical). (B) Diagrams showing in blue the position of the L18pL5e domain in Arabidopsis uL18-like, *E. coli* uL18 and *S. oleracea* uL18c proteins (Left panel). Tri-dimensional structures depicting the 3 α helices and 3 β sheets of the L18pL5e RNA binding domain as present in the *E. coli* uL18 (PDB ID: 4YBB) and the *S. oleracea* plastid uL18c (PDB ID: 5XBP). The structural models of uL18-like proteins were generated with Phyre2 (Right panel). Beta sheet are shown in yellow, alpha helices in red.

**Figure 2:**
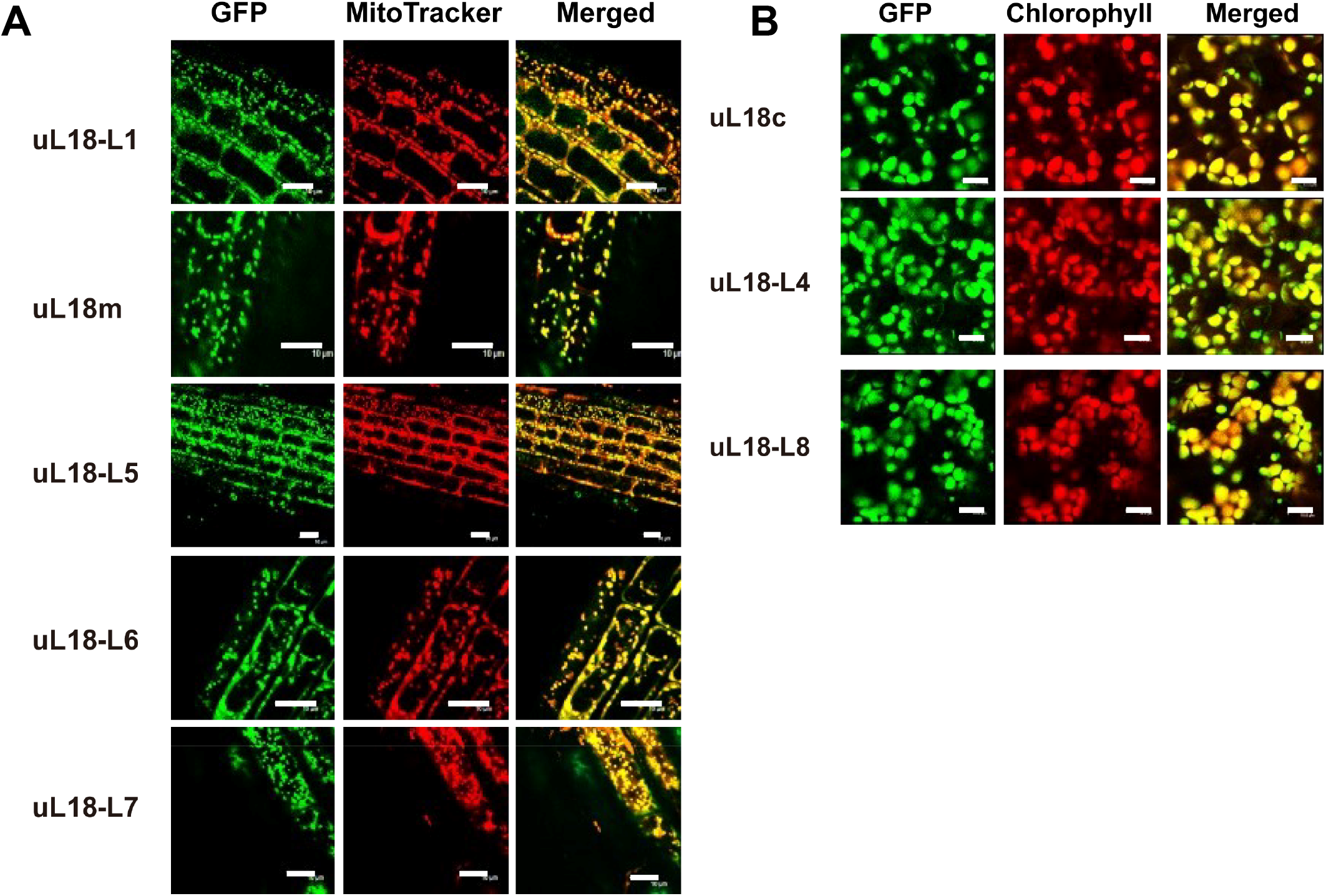
The Arabidopsis uL18 and uL18-like proteins are transported into mitochondria or plastids. Confocal microscope images showing the subcellular distribution of GFP translational fusions involving the indicated uL18 and uL18-like proteins. (A) Pictures taken from root cells of young Arabidopsis transgenic plants. Left: GFP fluorescence, center: Mitotracker Red marker, right: merged signals. (B) Pictures taken from leaf cells of Arabidopsis transgenic plants. Left: GFP fluorescence, center: chlorophyll fluorescence, right: merged signals. White scale bars correspond to 10 μm.

### *ul18-l1* mutant plants have a strongly retarded growth phenotype

To get insight into the function of uL18-Like proteins, we identified a first mutant in the *uL18-L1* gene (Figure 3A). Under conventional greenhouse condition, homozygous *ul18-l1* plants displayed a marked slow growth phenotype and harboured small twisted leaves (Figure 3B). The mutant plants were late flowering and produced seed with lower germination capacity compared to the wild type (Figure 3B). All these phenotypes could be complemented by the expression of an HA-tagged copy of uL18-L1 in the mutant, supporting that the observed alterations were due to the inactivation of AT1G08845 (Figure 3B). The mitochondrial localization of uL18-L1 was then confirmed by probing total, plastid and mitochondrial protein fractions prepared complemented plants (Supplementary Figure S2). Altogether, the results demonstrated the strict mitochondrial localization of the uL18-L1 protein.

**Figure 3:**
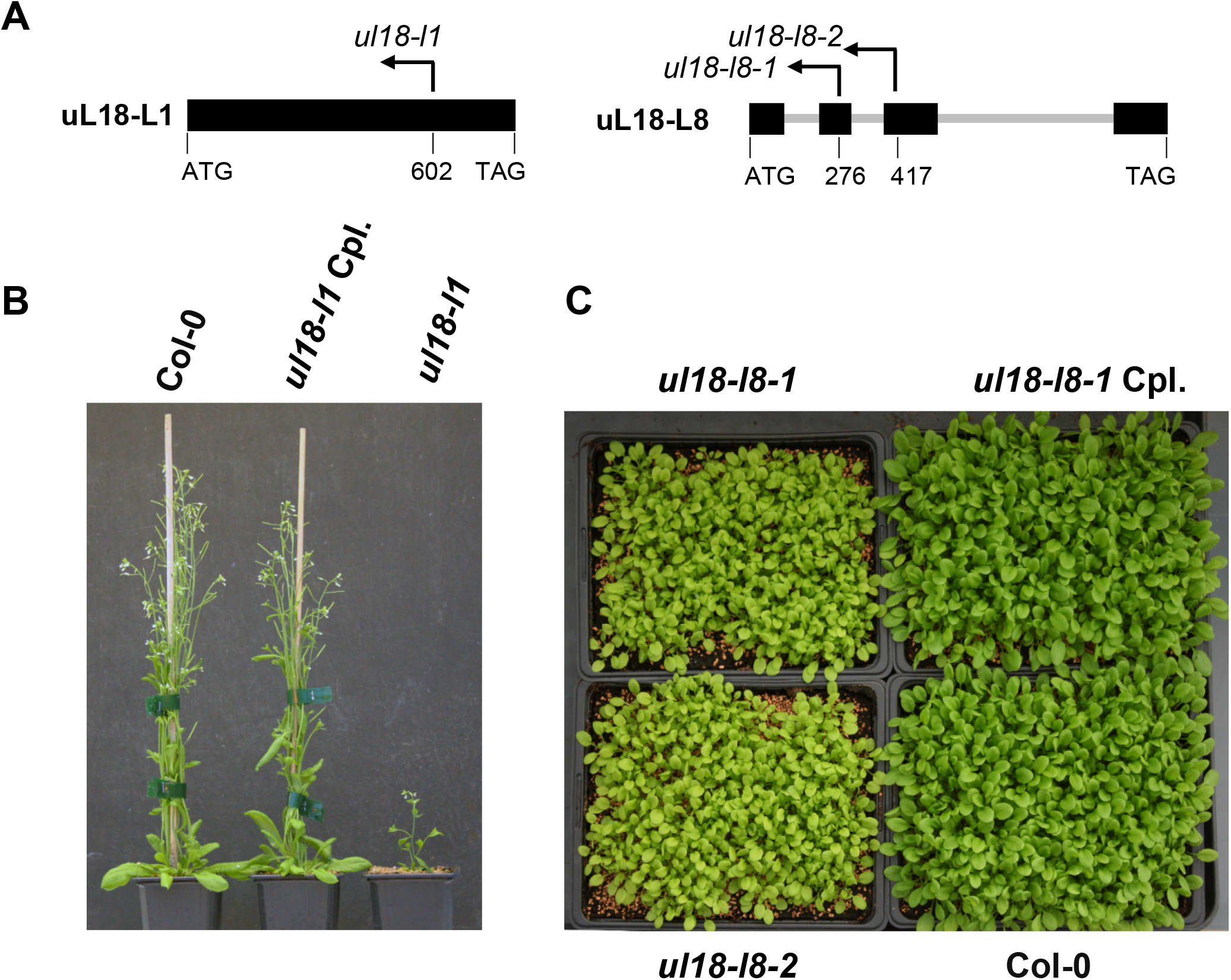
*ul18-l1* and *uL18-l8* mutants display altered developmental phenotypes. (A) Schematic representations of uL18-L1 and uL18-L8 gene structures with exons shown as black boxes, introns as thick gray lines. Arrows materialized the positions of T-DNA insertions and corresponding mutant alleles are indicated. (B) Photograph of 8-week-old plants showing the reduced size of *ul18-l1* homozygous mutants compared with wild-type and genetically complemented (Cpl) plants. (C) Photograph of 4-week-old plants at the rosette stage showing the slightly reduced size and pale green color of *l18-l8* homozygous mutants compared with wild-type and genetically complemented (Cpl) plants.

### *ul18-l1* mutant plants show strongly reduced levels of respiratory complex I

The mitochondrial localization of the uL18-L1 protein strongly suggested that the developmental alterations of *ul18-l1* plants could result from altered respiration. We thus monitored the accumulation level of the different respiratory complexes in comparison with the wild type by blue-native gel analysis. Immunoblot analyses revealed trace amounts of fully assembled complex I in *ul18-l1* plants, an over-accumulation of complex IV and a 450 kDa complex I assembly intermediate in the mutant (Figure 4A). In-gel activity staining confirmed the marked reduction of respiratory complex I in *ul18-l1* plants, and an over-abundance of complex IV (Supplemental Figure S3), which correlated with increased steady state levels of Cox1, Cox2 and cytochrome *c* (Figure 4C). The reduction of complex I activity in the *ul18-l1* mutant was further confirmed by *in vitro* root growth assays which revealed a much weaker sensitivity of *ul18-l1* plants to rotenone compared to the wild type (Supplemental Figure S3B). In response, we could detect an overexpression of several components of the alternative respiratory pathway in *ul18-l1* plants, including the alternative mitochondrial NADH dehydrogenase *NDB4* and the alternative oxidase (Figure 4C and Supplemental Figure S4). These results indicated that uL18-L1 is crucial for the biogenesis and the activity of mitochondrial complex I.

**Figure 4:**
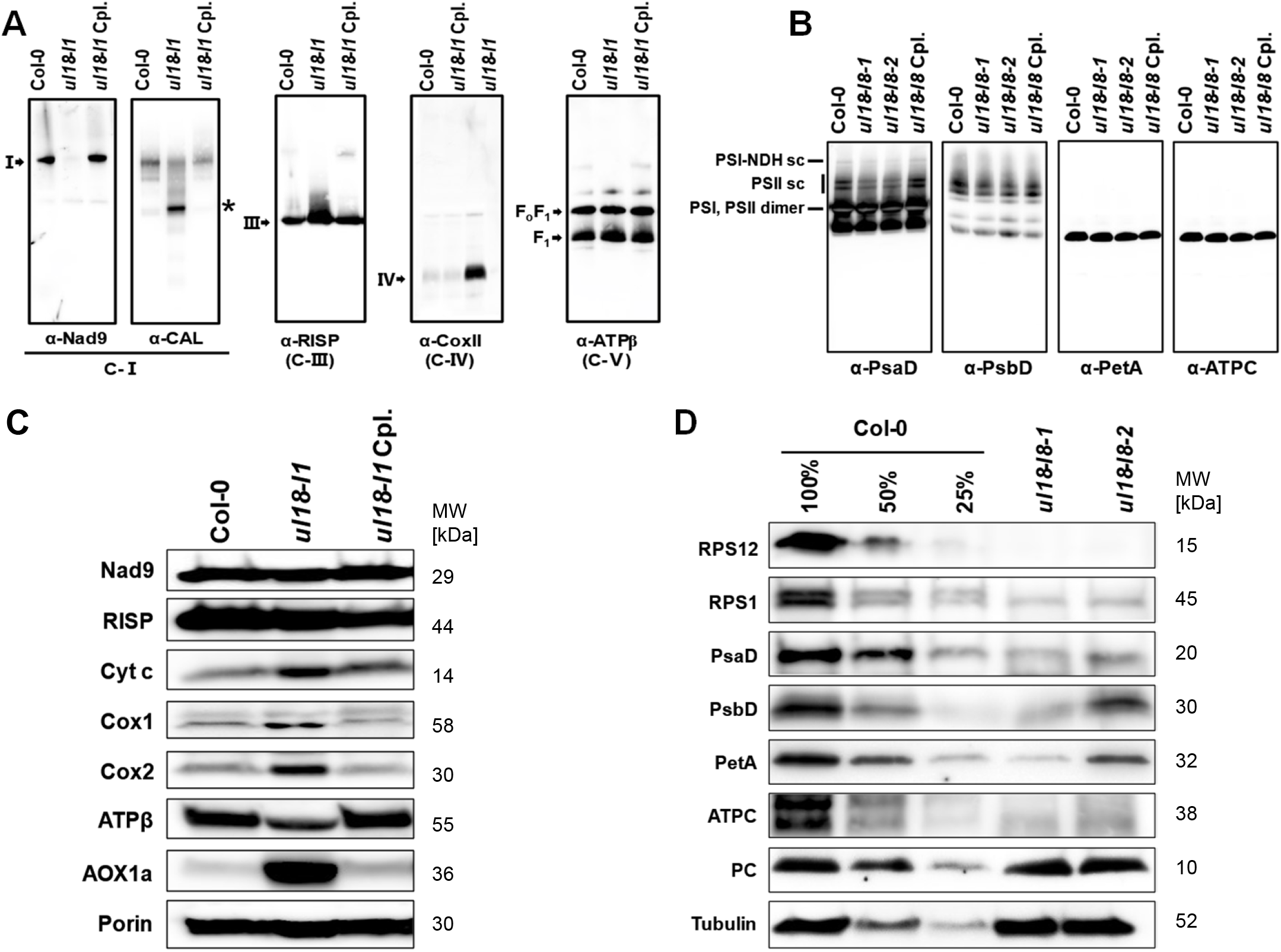
*ul18-l1* plants are defective in complex I and *ul18-l8* mutants under-accumulate plastid-encoded proteins. (A) Immunodetection of respiratory complexes in BN-PAGE blots made from mitochondrial extracts of wild-type (Col-0), *ul18-l1* mutant and complemented (Cpl) plants. Antibodies to NADH dehydrogenase subunit 9 (a-Nad9) and carbonate anhydrase-like (a-CAL) were used to detect fully assembled and assembly intermediates of complex I (C-I), respectively. Complex III (C-III) was detected with antibodies to the Rieske iron-sulfur protein (a-RISP), complex IV (C-IV) with anti-cytochrome oxidase subunit 2 antibodies (a-CoxII) and complex V (C-V) with ATP synthase subunit β antibodies (a-ATPβ). Respiratory complexes are designed by roman numerals, and the asterisk designates a complex I assembly intermediate. (B) Immunodetection of photosynthetic complexes on BN-PAGE blots made using thylakoid extracts from wild-type (Col-0), *ul18-l8* mutants and complemented (Cpl) plants. Antibodies to PsaB (a photosystem I (PSI) subunit), D2 (a photosystem II (PSII) subunit), PetA (a cytochrome *b_6_f* complex subuint) and ATPC (an ATP synthase subunit) were used. sc: super complex. (C) SDS PAGE immunoblots performed on total mitochondrial proteins extracts prepared from the indicated genotypes (see panel A for details) and probed with antibodies to subunits of respiratory complex I (Nad9), complex III (RISP), complex IV (Cox1 and Cox2), the ATP synthase (ATPβ), cytochrome *c* and the alterative oxidase (AOX). Porin was used as protein loading control. (D) Immunodetection of photosynthetic chain proteins in *ul18-l8* mutants compared to the wild type. Total plastid proteins purified from the indicated genotypes (see panel B for details) were probed with antibodies to subunits of the chloroplast ATP synthase (ATPC), photosystem II (PsbD), photosystem I (PsaD), the cytochrome *b_6_f* complex (PetA), ribosomal proteins (RPS12 and RPS1) and the plastocyanin. Tubulin was used as protein loading control. Dilution series of proteins extracted from wild type (Col-0) were used for signal comparison. Protein molecular weight (MW) is labeled on the right in kDa.

### The uL18-L1 protein is required for the splicing of two mitochondrial introns

Because of their homology to uL18, we suggested that uL18-type proteins may be RNA-binding proteins as well and that the uL18-L1 protein could therefore play a key role in the production of one or several mitochondria-encoded RNAs. To test this hypothesis, we determined the steady-state levels of precursor and mature mitochondrial transcripts by quantitative RT-PCR and calculated the splicing efficiency of mitochondrial introns in both wild-type and *ul18-l1* plants (Figure 5 and Supplemental Figure S5). A highly pronounced decrease in the splicing efficiency of *nad5* intron 4 could be detected as well a weak reduction for *nad2* intron 1 splicing in *ul18-l1* plants (Figure 5A). RNA gel blot confirmed these observations and indicated that no detectable amount of mature *nad5* mRNA could be found in the *ul18-l1* mutant (Figure 5D). The decrease in *nad2* intron 1 splicing was also verified but the production of mature *nad2* mRNA was much less affected than that of *nad5* and is therefore less likely to limit the production of complex I in *ul18-l1* plants (Figure 5 and Supplemental Figure S5). Altogether, these results indicated that the uL18-L1 protein is indispensable for the *cis*-splicing reaction involving *nad5* intron 4 and, to a lesser extent, also for the removal of *nad2* intron 1.

**Figure 5:**
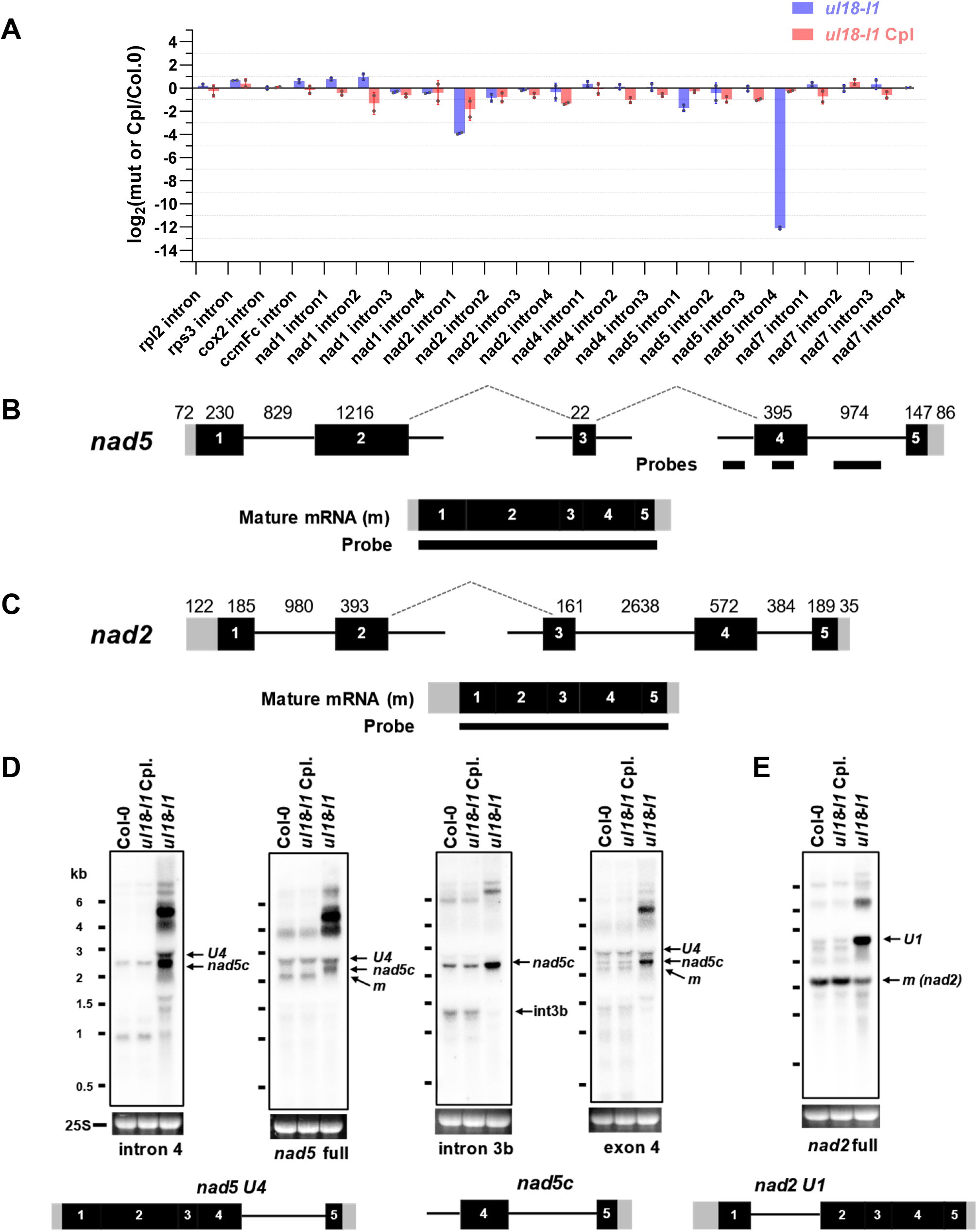
The *ul18-l1* mutant is impaired in *nad5* intron 4 and *nad2* intron 1 splicing. (A) Quantitative RT-PCR analysis measuring the splicing efficiencies of mitochondrial introns in *ul18-l1* and complemented (Cpl) plants. The bars depict log_2_ ratios of splicing efficiencies in *ul18-l1* and complemented plants to the wild type (Col-0). Three technical replicates and two independent biological repeats were used for each genotype. SD are indicated. (B) Schematic representation of the 3 *nad5* precursor transcripts which are fused by two *trans*-splicing events to produce the mature (m) *nad5* mRNA. Black boxes materialized exons. Introns are shown as thin lines, 5’ and 3’ UTRs as grey boxes. The nucleotide length of gene segments and probes used to interpret RNA gel blot results are also indicated. (C) Schematic representation of nad2 precursors and mature mRNA, using the same iconography as in (B). (D) RNA gel blot analysis showing the accumulation profiles of nad5 and nad2 transcripts in the ul18-l1 mutant compared to wild-type (Col-0) and complemented plants (Cpl). The blots were performed on total RNA prepared from flowers of the indicated genotypes. Used probes are indicated below hybridization results. Ethidium bromide staining of the 25s ribosomal RNA serves as a loading control. Schematic representations of nad5 U4, nad5c and nad2 U1 precursor transcripts are depicted below the blots to help interpreting the results. RNA marker sizes are indicated (kb). m, mature mRNAs.

### The *ul18-l8* mutant plants display altered photosynthesis

To further explore the functions of uL18-like proteins, we isolated two T-DNA insertion mutants in the *uL18-L8* gene whose encoded protein is transported into plastids (Figure 2B). Both mutant lines displayed a pale green phenotype associated with a slightly retarded growth (Figure 3C). These alterations could be reversed by expression an HA-tagged copy of uL18-L8 in one of the mutants, confirming that they were produced by the inactivation of *uL18-L8*. Pigment composition revealed that both mutants have a slight decrease in chlorophyll and carotenoid contents (Supplemental Table S4). Chlorophyll fluorescence further showed that the maximum quantum efficiency of PSII (Fv/Fm) was decrease by around 11% in both allelic mutants (Supplemental Table S4) and that fraction of open PSII centres (qL) decreased by 40%. Moreover, immunoblots of subcellular fractions revealed that uL18-L8 is enriched in isolated chloroplasts and is detected solely in the stromal fraction (Supplementary Figure S2). Overall, this analysis demonstrated that the loss of the uL18-L8 protein has a moderate but significant impact on the light energy efficiency of photosynthesis.

### The uL18-L8 protein is essential for the splicing of the first intron of *rps12* in plastid

To get insights into the function of L18-L8, quantitative RT-PCR were performed to comparatively measure the steady-state level of plastid-encoded transcripts in *ul18-l8* plants. We first observed that no plastid transcript suffered from a global destabilization in any of the mutants (Supplemental Figure S6). However, an around 64-fold decrease in the splicing efficiency of the first intron of the *rps12* transcript could be detected (Figure 6A). The mature *rps12* ORF is contained in a di-cistronic transcript along with the *rps7* gene and is produced *via* a *trans*-splicing reaction involving a first precursor comprising the first exon of *rps12* plus *rpl20* and a second precursor carrying the last two *rps12* exons as well as the *rps7* gene (Figure 6B). RNA gel blot analysis confirmed that no mature *rps12* di-cistronic mRNA could be detected in *ul18-l8* plants and that both *rps12* precursor transcripts over-accumulated compared to the wild type (Figure 6C). The splicing efficiency of introns contained in plastid tRNAs was then evaluated by RNA blot analysis and did not show any reduction (Supplemental Figure S7A). In contrast to plastid-encoded ribosomal RNAs (Supplemental Figure S7C), most mature plastid tRNAs over-accumulated in both mutants (Supplemental Figure S7A). Immunoblots of leaf extracts revealed that no trace of RPS12 could be detected in any of the *l18-l8* mutants (Figure 4D), correlating with the strong reduction of *rps12* intron splicing detected in these plants (Figure 6). A strong reduction in the steady state levels of all tested plastid-encoded proteins (RPS12, RPS1, PsaD, PsbD, PetA and ATPc) was also observed in *l18-l8* plants. These proteins accumulated at around 25% of wild-type levels, although the decrease appeared less pronounced in the *l18-l8-2* mutant line. Conversely, the level of the nuclearly-encoded plastocyanin was found unchanged in the mutants (Figure 4D). The analysis of photosynthetic complexes indicated that the Cytb6*/f* and ATP synthase complexes accumulated to the same level in wild-type and mutant plants, whilst the level of PSII complex was slightly reduced in *ul18-l8* mutants (Figure 4B).

**Figure 6:**
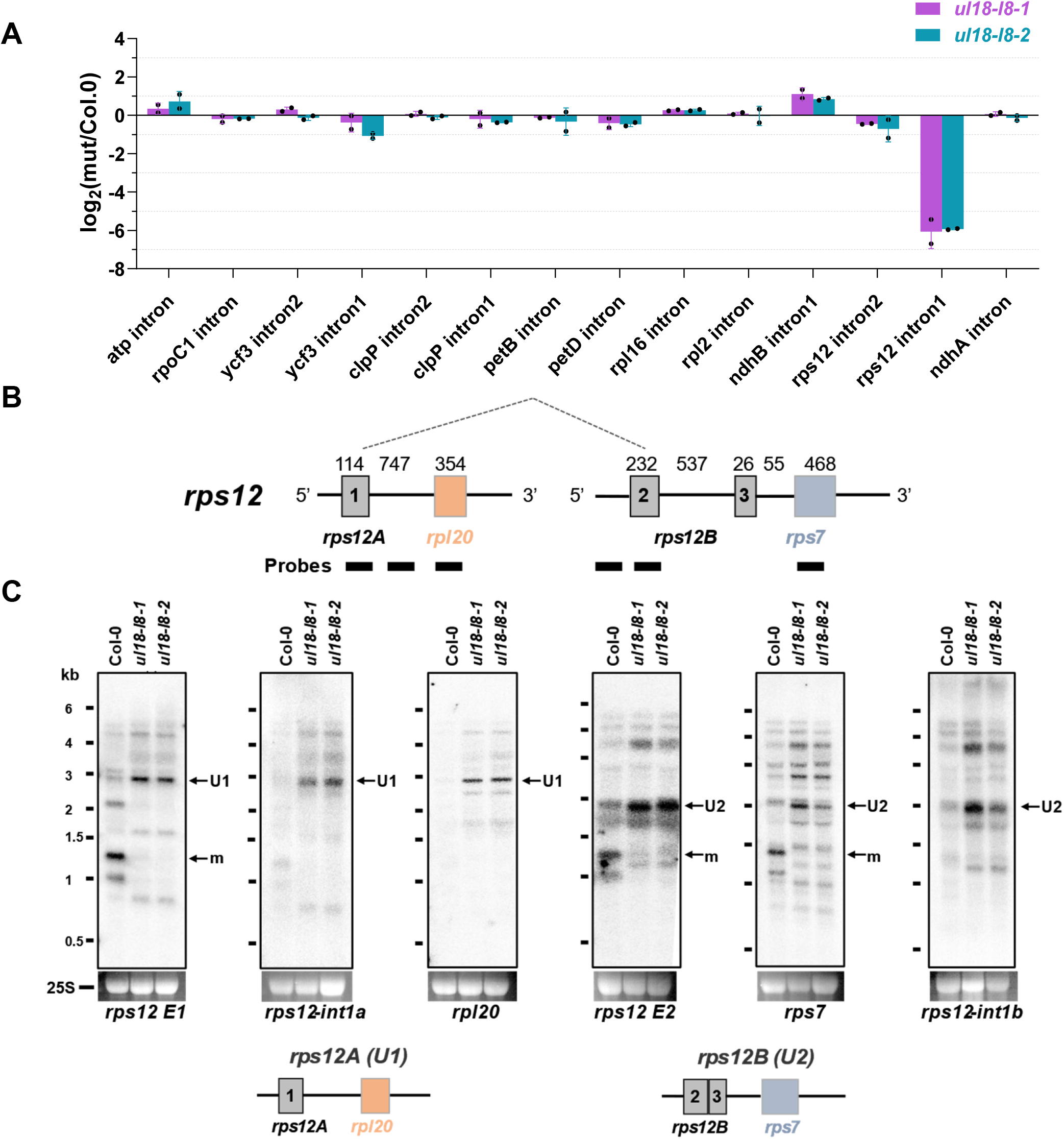
The *ul18-l8* mutants show a strong reduction in *rps12* intron 1 splicing. (A) Quantitative RT-PCR analysis measuring the splicing efficiencies of plastid introns in *ul18-l8* mutants. The bars show log2 ratios of splicing efficiencies in *ul18-l8* plants to the wild type (Col-0). Three technical replicates and two independent biological repeats were used for each genotype; standard errors are indicated. (B) Schematic representation of the two *rps12* precursor transcripts that are fused by *trans*-splicing to produce the mature (m) *rps12* mRNA. Black boxes materialized exons. Introns are shown as thin lines, 5’ and 3’ UTRs as grey boxes. The nucleotide length of gene segments and probes used to interpret RNA gel blot results are shown. (C) RNA gel blot analyses showing the accumulation patterns of *rps12* transcripts in *ul18-l8* mutants in comparison with the wild type. The blots were performed on total RNA prepared from 4-week-old leaves of the indicated genotypes. Used probes are specified below each blot. Ethidium bromide staining of the 25s ribosomal RNA serves as a loading control. Schematic representations of unspliced *rps12A* (U1) and *rps12B* (U2) precursor transcripts are depicted below the blots. RNA marker sizes are indicated (kb). m, mature mRNA.

Overall, these results demonstrated that the uL18-L8 protein is indispensable for the *trans*-splicing reaction involving the first intron of *rps12* in plastids and that *l18-l8* mutants consequently produce highly reduced levels of the RPS12 ribosomal protein resulting in a significant decrease in the production of several plastid-encoded proteins.

### L18-L1 and uL18-L8 do no associate with mitochondrial or plastid ribosomes

To see whether the uL18-L1 and uL18-L8 proteins conserved a capacity to associate with mitochondrial or plastid ribosomes, the size of complexes engaging these two proteins was measured by density gradient sedimentation. Most previously identified mitochondrial or plastid group II intron splicing factors reside in high molecular weight ribonucleoprotein complexes that are, however, lighter than ribosomes (12, 35). Mitochondrial or stromal extracts prepared from *ul18-l1* and *ul18-l8*-*1* complemented plants were fractionated on sucrose gradient and the recovered fractions were analysed by immunoblot assays. Both uL18-Like proteins were found in particles that were much smaller than mito- or chloro-ribosomes (Figure 7). Interestingly, uL18-L8 was found in lighter fractions compared with the plastid splicing factor mTERF4 (36). Identical results were obtained with overexpressing Arabidopsis cell lines in which the detection of uL18-L1 and uL18-L8 is greatly facilitated compared to organelles purified from complemented plants (Supplementary Figure S8A). Both proteins could be immunoprecipitated from these transgenic lines but none of the tested anti-ribosome antibodies indicated co-enrichment with mitoribosomes for uL18-L1 or chlororibosomes for uL18-L8 (Supplementary Figure S8B).

**Figure 7:**
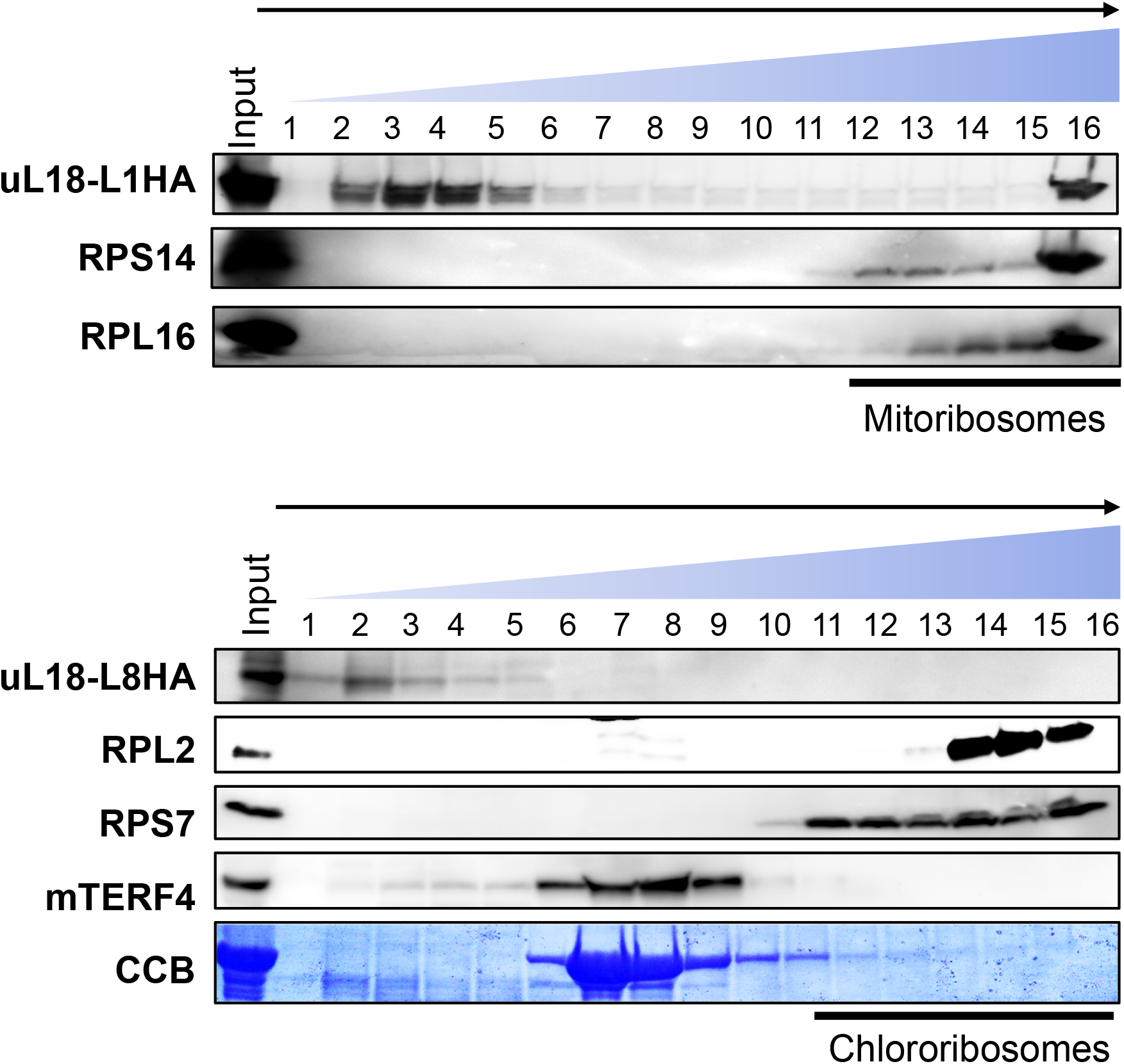
uL18-L1 and uL18-L8 do not take part in ribosomes. Mitochondrial (top) and stromal (bottom) extracts prepared from *ul18-l1* and *ul18-l8-2* complemented plants respectively were fractionated on 10-30% sucrose gradients. An equal volume of each recovered fraction was analyzed by protein gel blot with the indicated antibodies. Mitochondrial RPL16 and RPS14 as well as plastid RPL2 and RPS7 were used to localize mitochondrial or plastid ribosomes along the gradients. mTERF4 is a protein required for the splicing of multiple plastid group II introns (Hammani and Barkan, 2014). The αHA antibody detects uL18-L1-3HA and uL18-L8-3HA fusion proteins. The Coomassie blue staining of the membrane used for detecting the large subunit of Rubisco (RbcL) is shown below the hybridization results.

### L18-L1 and uL18-L8 associate with introns whose splicing they facilitate *in vivo*

To identify intron RNAs that associate with L18-L1 and uL18-L8 *in vivo*, both proteins were immunoprecipitated from over-expressing Arabidopsis cell lines with HA antibody and co-enriched RNAs were purified from the coimmunoprecipitates. Obtained RNAs were then used for cDNA synthesis and analysed by qPCR using primer pairs amplifying mitochondrial or plastid precursor mRNAs. Enrichment ratios were calculated in comparison to control immunoprecipitations performed with IgG-Protein A magnetic beads. We observed a very high and specific enrichment of uL18-L1 with *nad5* intron 4 and uL18-L8 with *rps12* intron 1 (Figure 8). Interestingly, *nad2* intron 1 was not co-enriched in uL18-L1 immunoprecipitates, suggesting that *nad2* intron 1 splicing defect detected in *ul18-l1* plants could be a secondary effect of complex I deficiency as observed in several independent Arabidopsis complex I mutants (26, 37–41). Both uL18-type proteins thus associate with their genetic target *in vivo*, supporting a direct role in the splicing of the concerned introns.

**Figure 8:**
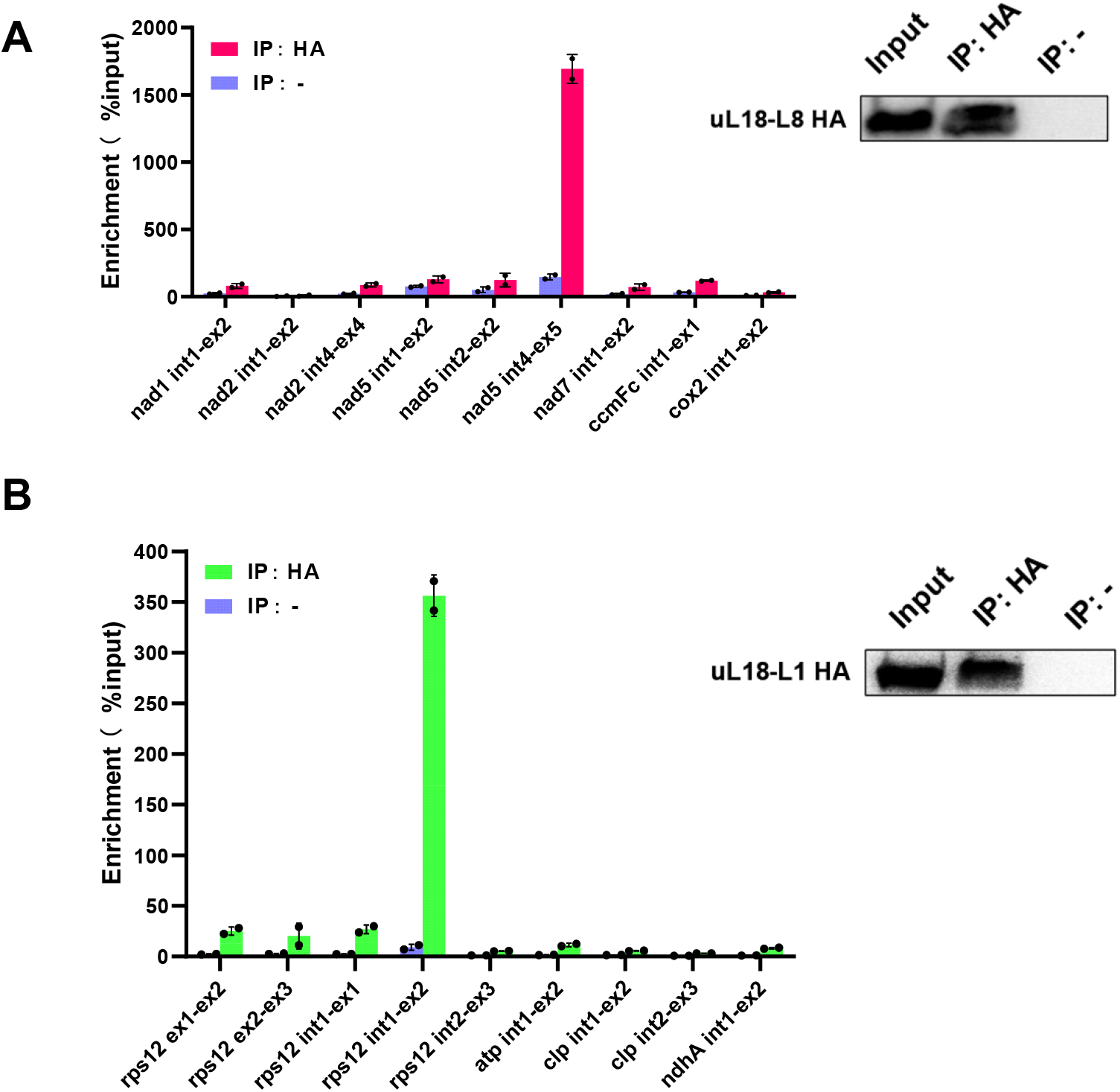
uL18-L1 and uL18-L8 specifically associate with *nad5* intron 4 and *rps12* intron 1, respectively. (A) Analysis of RNAs that coimmunoprecipitate with uL18-L1-3HA. Total extracts from transgenic Arabidopsis cell lines expressing uL18-L1-3HA were used for immunoprecipitation with anti-HA antibody (IP+) and IgG-Protein A magnetic beads (IP−) as negative control. RNAs were extracted from immunoprecipitation pellets and then analyzed by RT-qPCR using primer pairs positioned across intron/exon junctions of the indicated mitochondrial transcripts. Enrichment ratios calculated with the delta Ct method are shown. (B) RT-qPCR analysis of RNAs coimmunoprecipitating with uL18-L8-3HA. Similar experiment as in (A) expect that cells expressing uL18-L8-3HA were used as starting material and that primers pairs amplifying plastid transcript precursors were employed to analyze co-immunoprecipitated RNAs. Immunoblot results of input and immunoprecipitated fractions using the HA antibody (IP+) or IgG-Protein A magnetic beads (IP−) are shown.

## Discussion

### The uL18 RNA binding domain was recruited early in the evolution of terrestrial plants to produce intron-specific splicing factors for organelles

Ribosomal proteins (RP) are essential components of ribosomes that help structuring rRNAs to allow them to function optimally in protein synthesis. In recent years, several lines of evidence have documented that RPs can carry ribosome-independent functions (16, 42, 43). In this report, we revealed that two Arabidopsis proteins of the uL18 family play essential roles in group II intron splicing in chloroplasts or mitochondria. The uL18-L1 protein was found to be indispensable for the splicing of last intron contained in *nad5* pre-mRNAs in mitochondria, whereas the plastid uL18-L8 is required for the removal of *rps12* intron 1 (Figure 9). These processes cannot be considered as extra-ribosomal functions *sensu stricto*, since both uL18-L1 and uL18-L8 proteins were not found to reside in mitochondrial or plastid ribosomes, respectively (Figure 7). Our results support nevertheless the capacity of ribosomal protein binding domains to accommodate new RNA targets, and reveal for the first time that this class of proteins can acquire functions in intron splicing. These observations add support to an important evolutionary paradigm of nature that uses existing genes for functional diversification. In the present cases, duplicated and subsequently divergent copies of *uL18m* or *uL18c* were likely selected to play roles in organellar intron splicing at some point in the evolution of plants. The search for uL18 homologs from bacteria to seed plants indicates that the diversification of the uL18 family occurred early in vascular plant evolution. In bacteria or algae, a single *uL18* gene can be generally recognized, which encodes a protein sharing strong homologies with the ribosomal protein uL18c (Supplementary Figure S1). Non-vascular plants encode a few more uL18 like proteins whose closest homologs correspond to the uL18m, uL18c and uL18-L4 proteins. Gymnosperms and angiosperms encode similar corteges of uL18 and uL18-like proteins as in Arabidopsis, with the intriguing exception of uL18-6 that is specific to dicotyledonous plants (Supplementary Figure S1). It thus appears that the expansion of the uL18 family was initiated in non-vascular basal plants and that the first diverging uL18 member may have been the *uL18-L4* gene. These observations nicely corroborate previous analyses indicating that deviation of organellar introns from the classic structure of group II introns has occurred early in the evolution of terrestrial plants (44–46).

**Figure 9:**
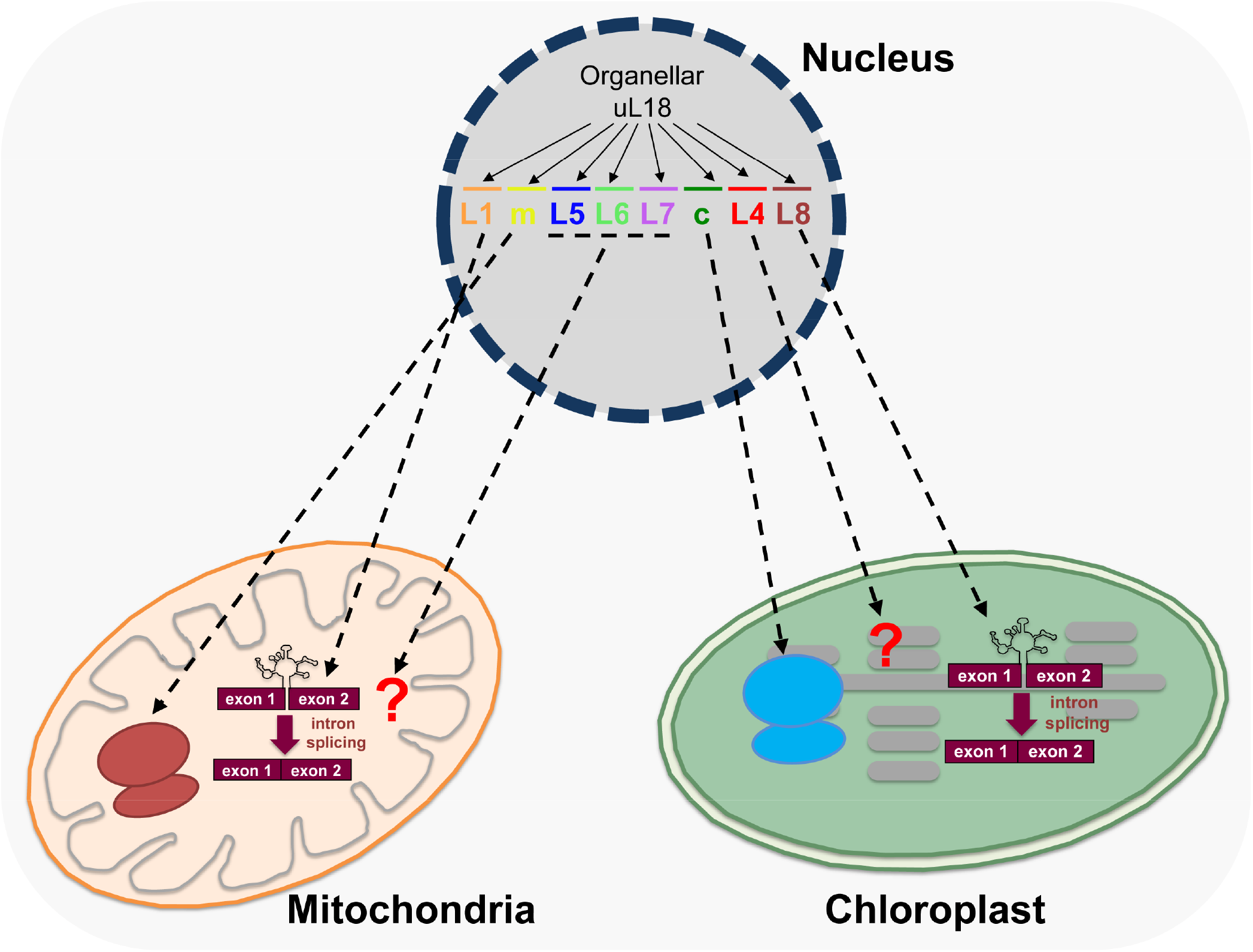
Graphical summary depicting the functional diversification of the uL18 ribosomal protein family in Angiosperms. The uL18 protein family comprises 8 members that evolved from ancestral organellar *uL18* genes. Five members are transported into mitochondria and 3 into plastids. Two uL18 proteins (uL18m and uL18c) have retained the original uL18 function and are subunits of organellar ribosomes. Among the other six uL18-like members, uL18-L1 and uL18-L8 have evolved towards splicing and specifically associate *in vivo* with introns whose splicing they promote.

Organellar introns derive from bacterial counterparts and can be classified as group I or group II based on specific structural characteristics. Introns contained in vascular plants organellar genomes are mostly of type II and fold into a conserved structure comprising six helical domains (DI–DVI) that radiate from a central hub (47). Autonomous group II introns encode a maturase that is essential for intron mobility and splicing. Most organellar introns in seed plants show degenerate structures compared to canonical group II introns or lack structural features that are important for removal (45, 46). Unlike in bacteria, these deviant group II introns are maintained in plant organellar genomes as splicing-only elements. They may be mobilized *via* unusual splicing mechanisms, but a tolerance to structural changes permitted by the selection of a compensatory accessory machinery is an evolutionary strategy that is chosen to maintain efficient splicing of derivatized introns in organelles (46, 48). Various kinds of RNA binding proteins have been recruited to play roles in organellar intron splicing like aminoacyl tRNA synthetases, a peptidyl-tRNA hydrolase, pseudouridine synthase homologs or proteins derived from a bacterial ribosome assembly factors (11, 49–52). Minor evolutionary tinkering seems thus sufficient to confer novel activities in intron splicing to these RNA binding proteins, like for the uL18 homologs. By associating with their target intron, these factors form ribonucleoprotein complexes that could help intron folding and stabilize their structure in a catalytically active form (53). However, advanced biochemical and structural analyses are necessary to understand how uL18-Like proteins and other proteinaceous factors facilitate group II intron splicing in plant organelles.

### The uL18-L1 and uL18-L8 proteins could associate with 5S rRNA-like structures in their target intron

In most cases, the evolutionary recruitment of RPs towards new sites is permitted because the structure of newly targeted mRNA regions resembles that of rRNA domains to which they associate in the ribosomes (54–61). The uL18 protein is the most important factor for incorporating the 5S rRNA into the 50S ribosomal subunit (62). The 5S rRNA is a small RNA of around 120 nucleotides long and adopts a conserved tertiary structure comprising various helices as well as terminal and internal loops (63). The most important structural determinants for uL18 anchoring are contained in its β domain comprising two helices plus one internal and a terminal loop (31). Upon binding, the uL18 protein increases the stability of 5S rRNA, modifies its structure and enhances its affinity of uL5 (64, 65). Our structural predictions showed that C-terminal regions of uL18-L1 and uL18-L8 adopt the same structure as ribosomal uL18s and that the three β sheets forming the RNA binding surface of uL18 are also found in the predicted structures of these two uL18 like proteins (Figure 1). This strongly suggests that the recruitment of uL18-like proteins in splicing was permitted because regions within *nad5* intron 4 and *rps12* intron 1 likely resemble the β domain of 5S rRNA, at least enough so that uL18-L1 and uL18-L8 proteins could associate with them in a productive way. Interestingly, the recent structures of mammalian mitoribosomes showed that the uL18 binding domain can accommodate different but structurally similar RNA targets since tRNAs serve as architectural replacements for the 5S rRNA in these ribosomes (66, 67). Therefore, the search for regions within *nad5* intron 4 and *rps12* intron 1 folding similarly to the β domain of 5S rRNA may reveal the binding sites of uL18-L1 and uL18-L8. As uL18 does with 5S rRNA, the binding of uL18-L1 and uL18-L8 could induce local structural changes to the RNA regions to which they bind and thereby help their target intron to fold in an active form. These structural modifications may also permit the recruitment of other protein factor(s) necessary for splicing, similarly to the impact that the bacterial uL18 has on increasing the uL5 binding affinity for 5S rRNA. Finding the exact binding sites of uL18-L1 and uL18-L8 within their target introns represents an important challenge to better understand the role of uL18-L1 and uL18-L8 in group II intron splicing. Deciphering the functions of other uL18-like proteins will reveal whether they also play roles in organellar intron splicing or whether they are involved in other RNA expression steps in plant organelles. Together, these results will better document how ribosomal proteins can evolve towards new functions in RNA metabolism and how they can adapt to associate and process new RNA targets.

## Supplementary data

Supplementary Data are available online.

## ACKNOWLEDGMENTS

ANR MITRA (ANR-16-CE11-0024-01) to H. M. China Scholarship Council to C. W. The IJPB benefits from the support of Saclay Plant Sciences-SPS (ANR-17-EUR-0007). This work has benefited from the support of IJPB’s Plant Observatory technological platforms.

---

**Supplemental Figure S1:**
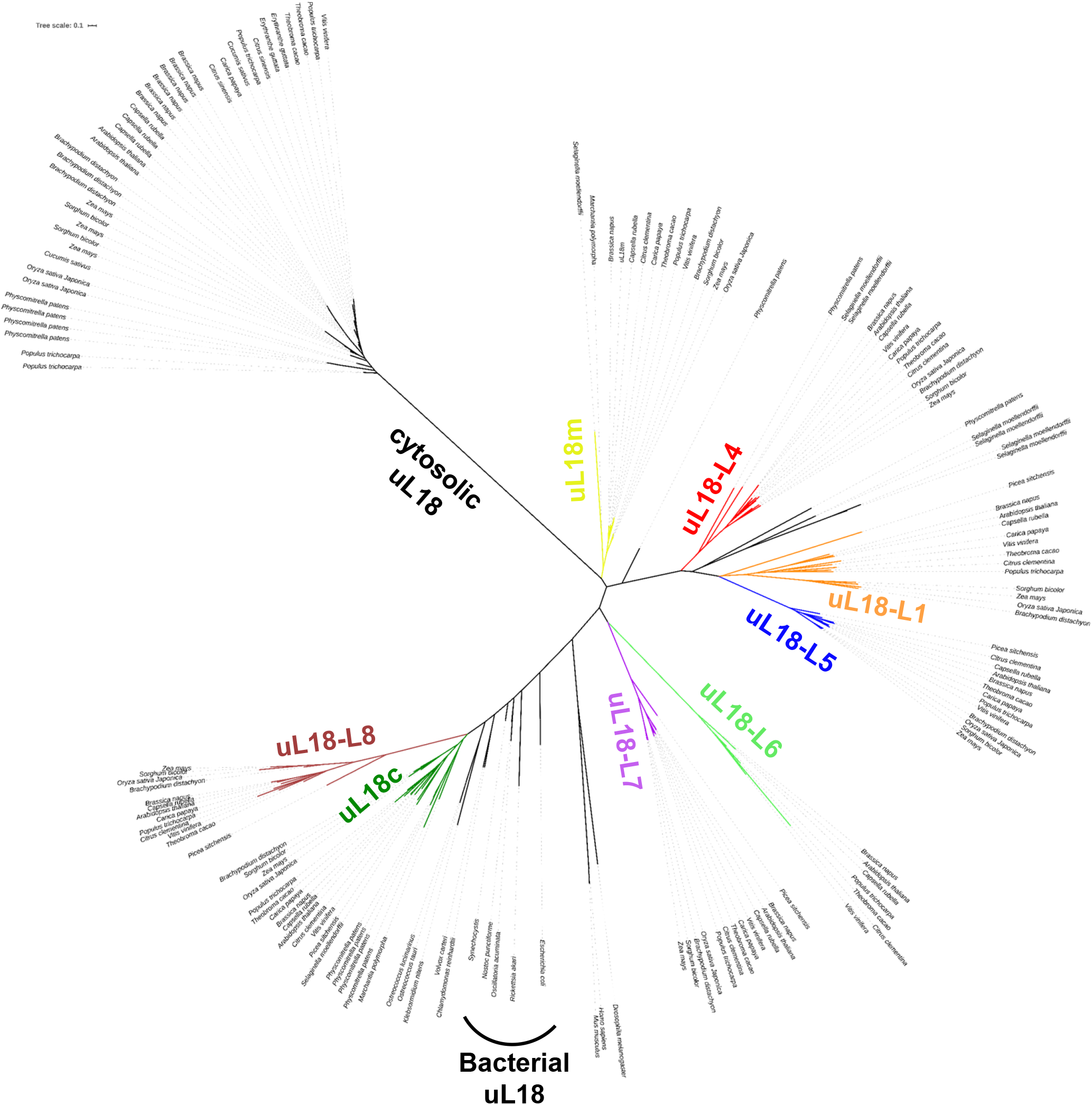
The uL18 and uL18-like proteins are conserved in monocots and dicots. Sequence distance neighbor-joining tree of uL18 or uL18-like protein homologs from a selected panel of bacteria, animalia, Chlorophyta, bryophytes, gymnosperms, dicotyledons and monocotyledons. All accession numbers of the proteins reported here are listed in Supplemental Table S3. These alignments reveal likely orthologs for each Arabidopsis uL18-like in monocots and dicots, supporting their functional conservation in terrestrial plants. Sequence names are either GenBank accession numbers or Genome Initiative locus names. The tree was generated using MEGA7 (www.megasoftware.net) and visualized via iTOL (Interactive Tree Of Life, https://itol.embl.de/).

**Supplemental Figure S2:**
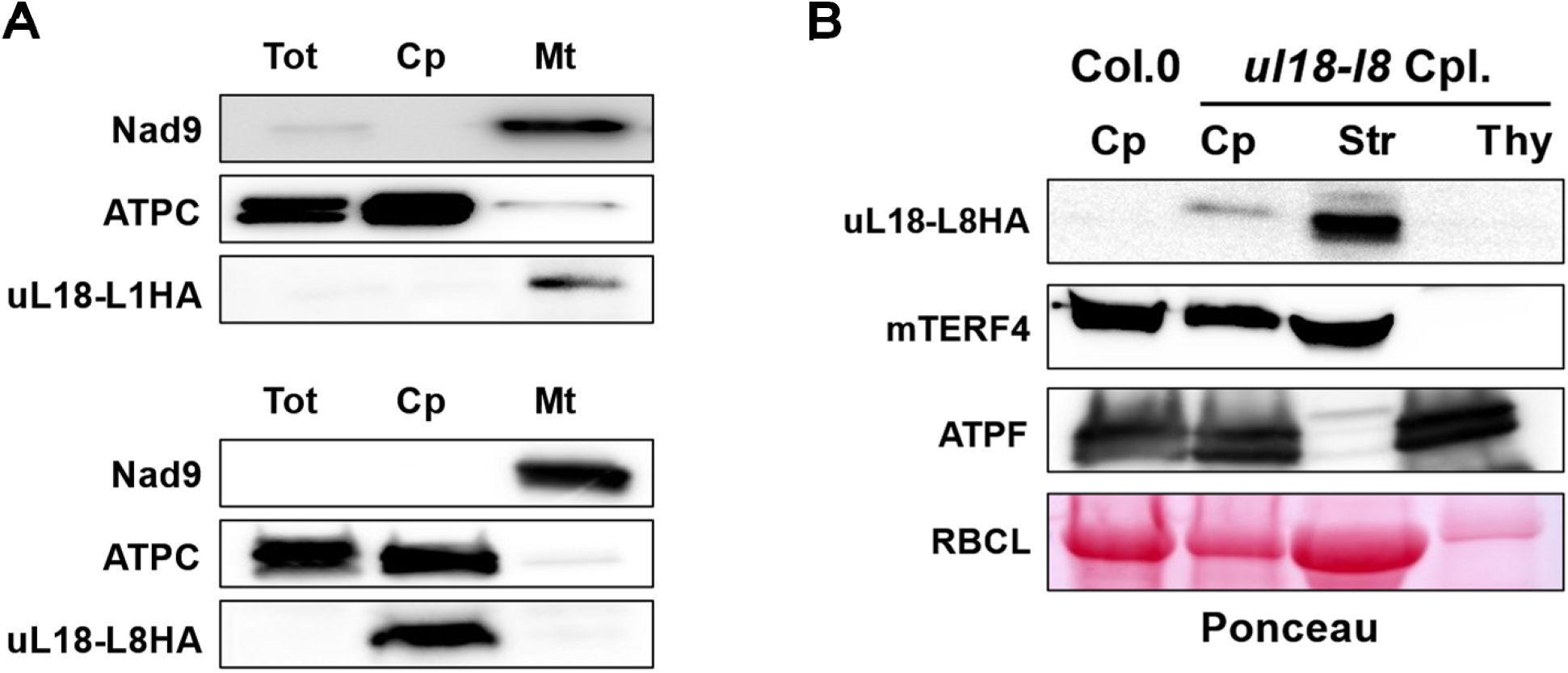
Subcellular distributions of uL18-L1 and uL18-L8 proteins. (A) Total (Tot), chloroplast (Cp) and mitochondrial (Mt) protein extracts were prepared from genetically complemented *ul18-l1* and *ul18-l8-2* lines and analyzed by immunoblot assays with the indicated antibodies. The detection of the plastid ATP synthase subunit C (ATPC) and the mitochondrial NADH dehydrogenase subunit 9 (Nad9) were used to control the purity of plastid and mitochondrial extracts, respectively. The anti-HA monoclonal antibody was used to detect the uL18-L1-HA and uL18-L8-HA fusion proteins. (B) Detection of the uL18-L8 protein in chloroplast (Cp), stromal (Str) and thylakoid (Thy) fractions by immunoblot analysis. Antibodies to mTERF4 and ATPF were used as control for stromal and thylakoid fractions, respectively.

**Supplemental Figure S3:**
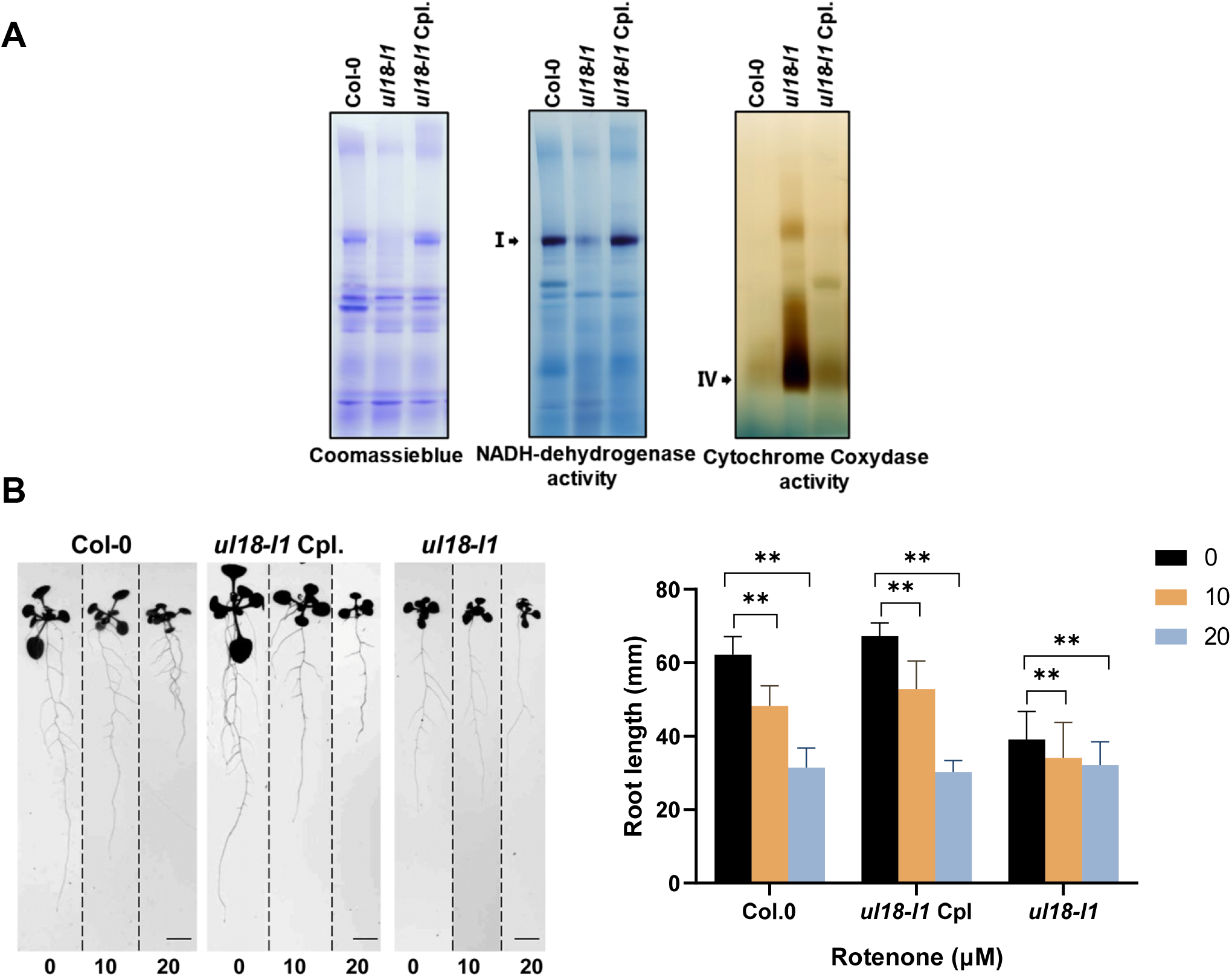
*ul18-l1* plants are complex I mutants and *ul18-l8* mutants show reduced photosynthetic activity. (A) Mitochondrial protein extracts prepared from wild-type (Col-0), mutant (*ul18-l1*) and complemented plants (Cpl) were separated in BN-PAGE gels and then stained with Coomassie blue or by in gel activity staining to estimate complex I (NADH dehydrogenase) and complex IV (cytochrome *c* oxidase) accumulation levels in the different genotypes. The different respiratory complexes are designated by their roman numeral. (B) Primary root length of wild-type (Col-0), mutant (*ul18-l1*) and complemented plants (Cpl) measured in the absence or in the presence of 10 μM or 20 μM of the complex I inhibitor rotenone. Plants were grown on vertical squared plates for 14 (Col-0 and Cpl) or 24 (*ul18-l1*) days prior to measurements (left panel). The values (right panel) are means of three independent biological replicates (error bars indicate SD). Asterisks indicate statistically significant differences between the inhibitor-treated versus the no-inhibitor conditions, for each genotype (Student’s t test for paired samples, P, 0.01).

**Supplemental Figure S4:**
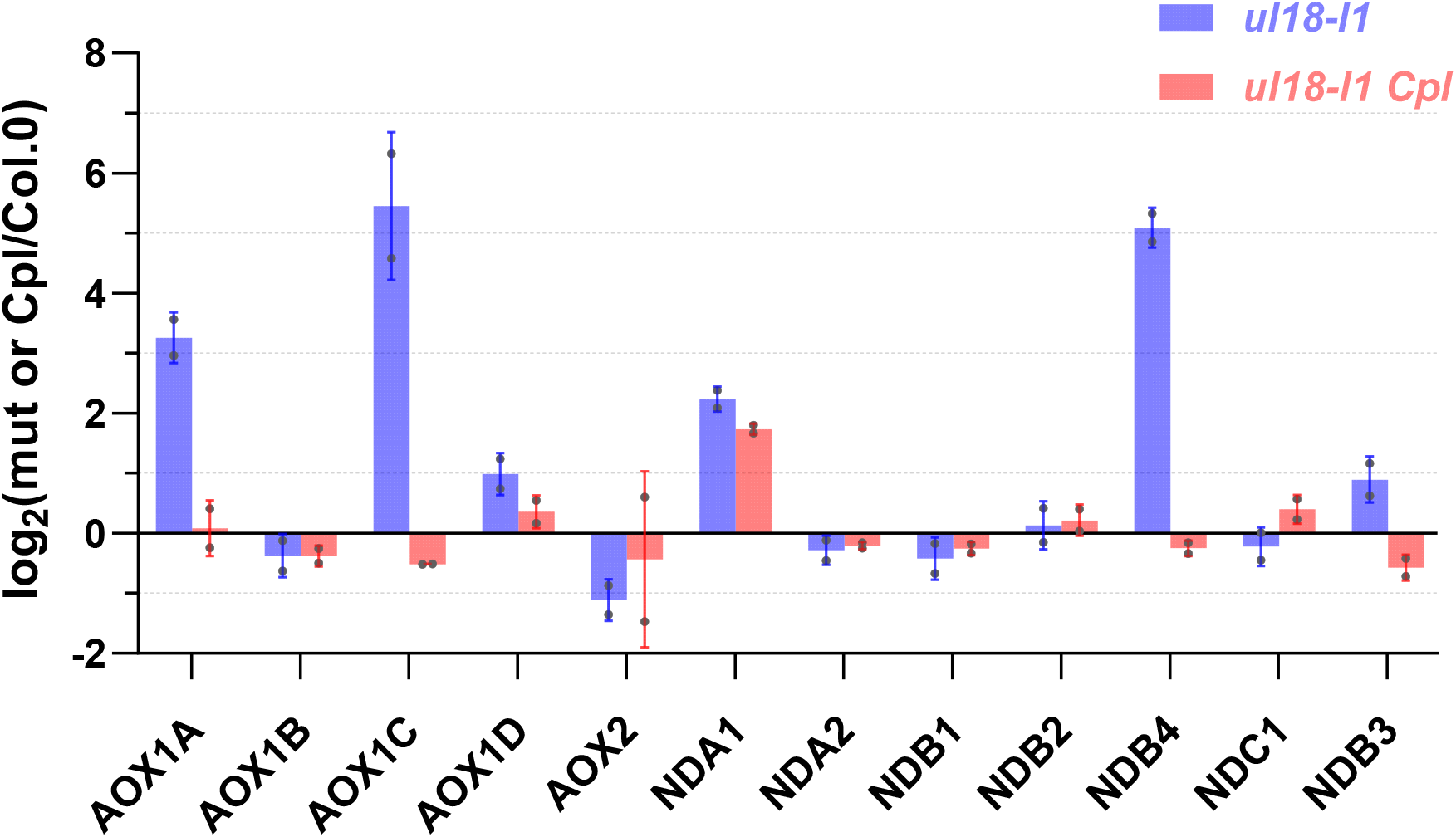
The alternative respiratory pathway is induced in *ul18-l1* mutant plants. Quantitative RT-PCR results measuring the relative accumulation levels of alternative oxidase (*AOX*) and NADH dehydrogenase (*NDA*, *NDB* and *NDC*) transcripts in *ul18-l1* mutant and complemented (Cpl) plants. Log_2_ ratios of mutant or Cpl to wild-type are shown. Two biological repeats and three technical repeats were performed for each genotype in this analysis.

**Supplemental Figure S5:**
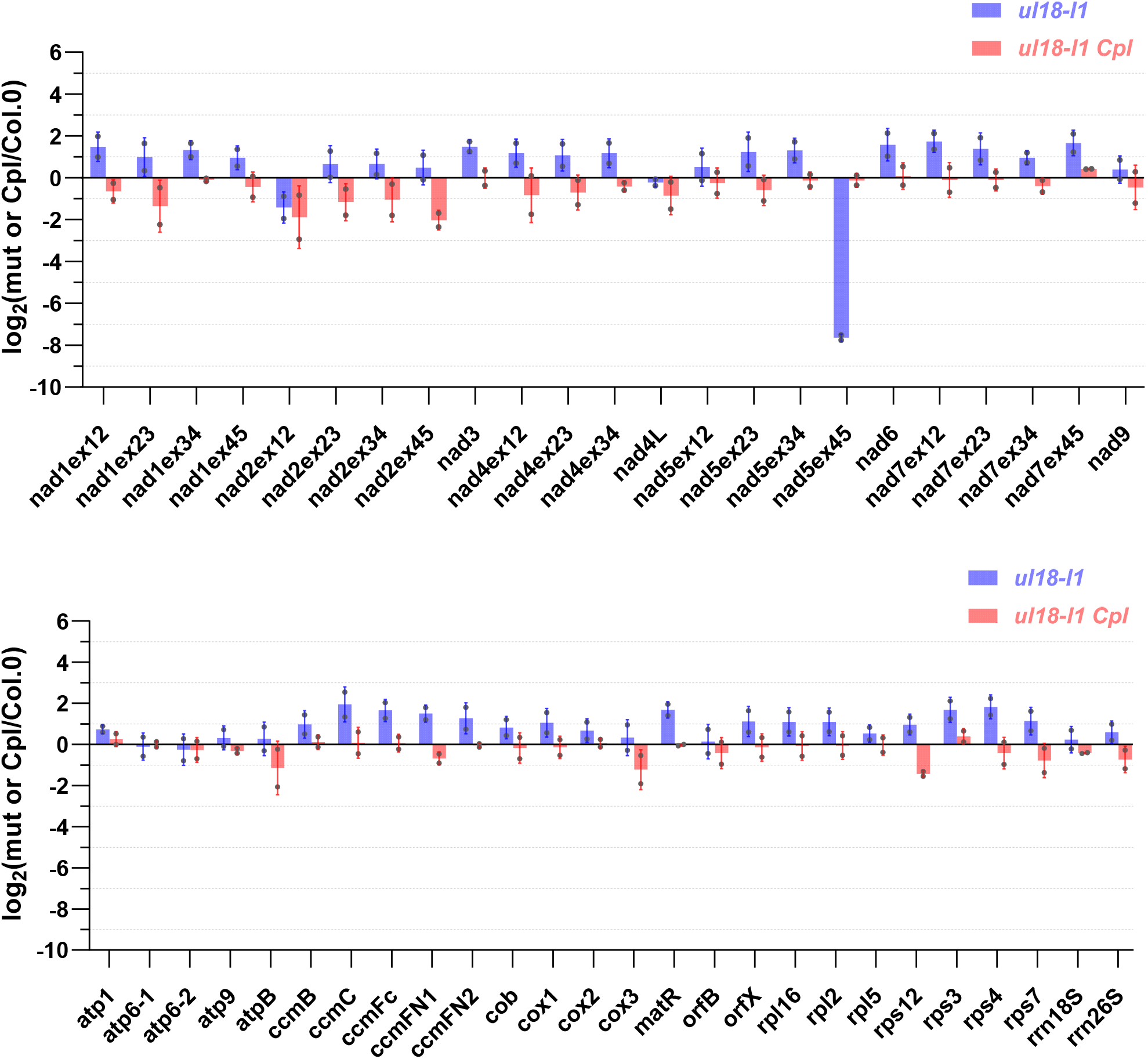
Quantitative RT-PCR analysis measuring the accumulation levels of mature mitochondria-encoded mRNAs in *ul18-l1* and complemented (Cpl) plants. A single PCR was performed for transcripts containing a sole exon. For intron-containing mRNAs, PCR were done using primer pairs across introns. The bars show log_2_ ratios of mature mRNA abundance in *ul18-l1* and complemented plants to the wild type (Col-0). Three technical replicates and two independent biological repeats were used for each genotype. Standard errors are indicated.

**Supplemental Figure S6:**
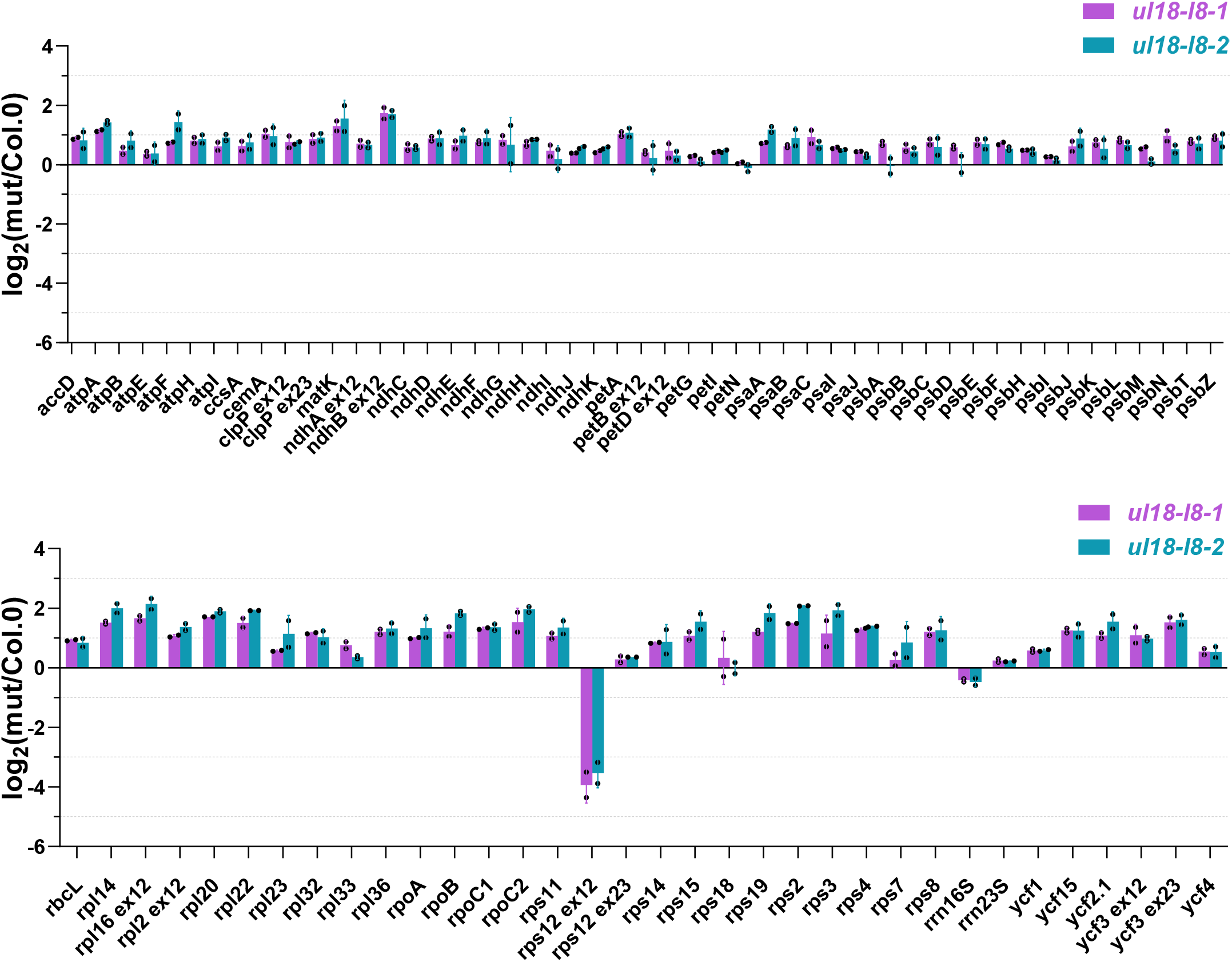
Quantitative RT-PCR analysis measuring the accumulation levels of mature plastid-encoded mRNAs in *ul18-l8* mutants. The graph depicts the log2 ratio of transcript levels in the mutants compared with those in wild-type. A striking decrease in the accumulation of *rps12* exon1-exon2 transcripts was observed, other transcripts showed insignificant reductions or weak increase. Three technical replicates and two independent biological repeats were used for each genotype. Standard errors are indicated.

**Supplemental Figure S7:**
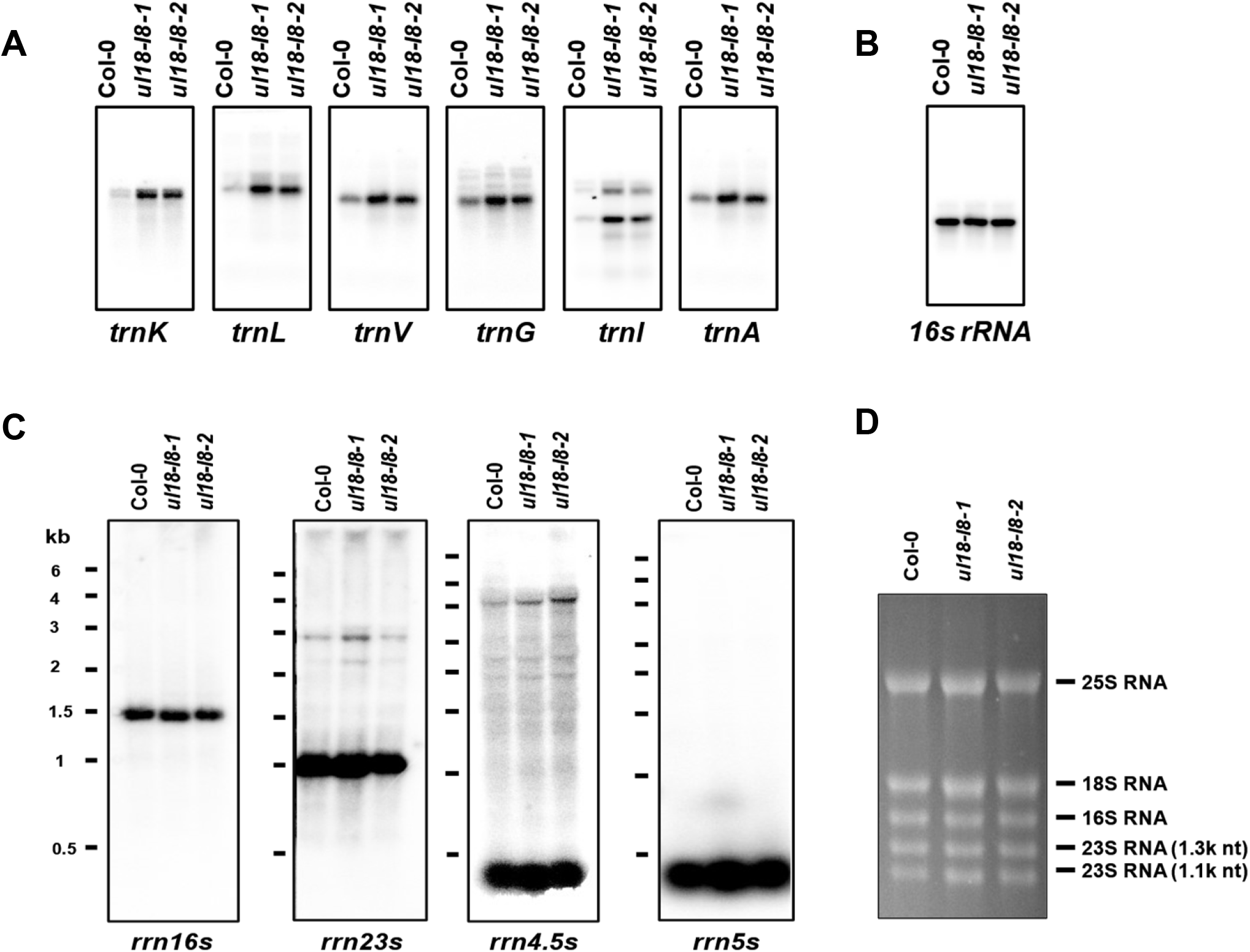
Plastid tRNAs and rRNA accumulation in *ul18-l8* mutants. The blots were performed on total RNA prepared from 4-week old rosette leaves of the indicated genotypes and used probes are mentioned below each blot. (A) RNA gel blot hybridizations showing the steady state levels of plastid tRNAs. (B) A blot identical to the ones shown in (A) and hybridized with a 16S rRNA probe was used as RNA loading control, since the expression level of this transcript is not unaffected in *ul18-l8* plants (see below). (C) RNA gel blot analyses measuring the steady state levels of plastid-encoded rRNAs. (D) Ethidium bromide staining of a representative formaldehyde denaturing gel used in (C) to prove equal loading between samples.

**Supplemental Figure S8:**
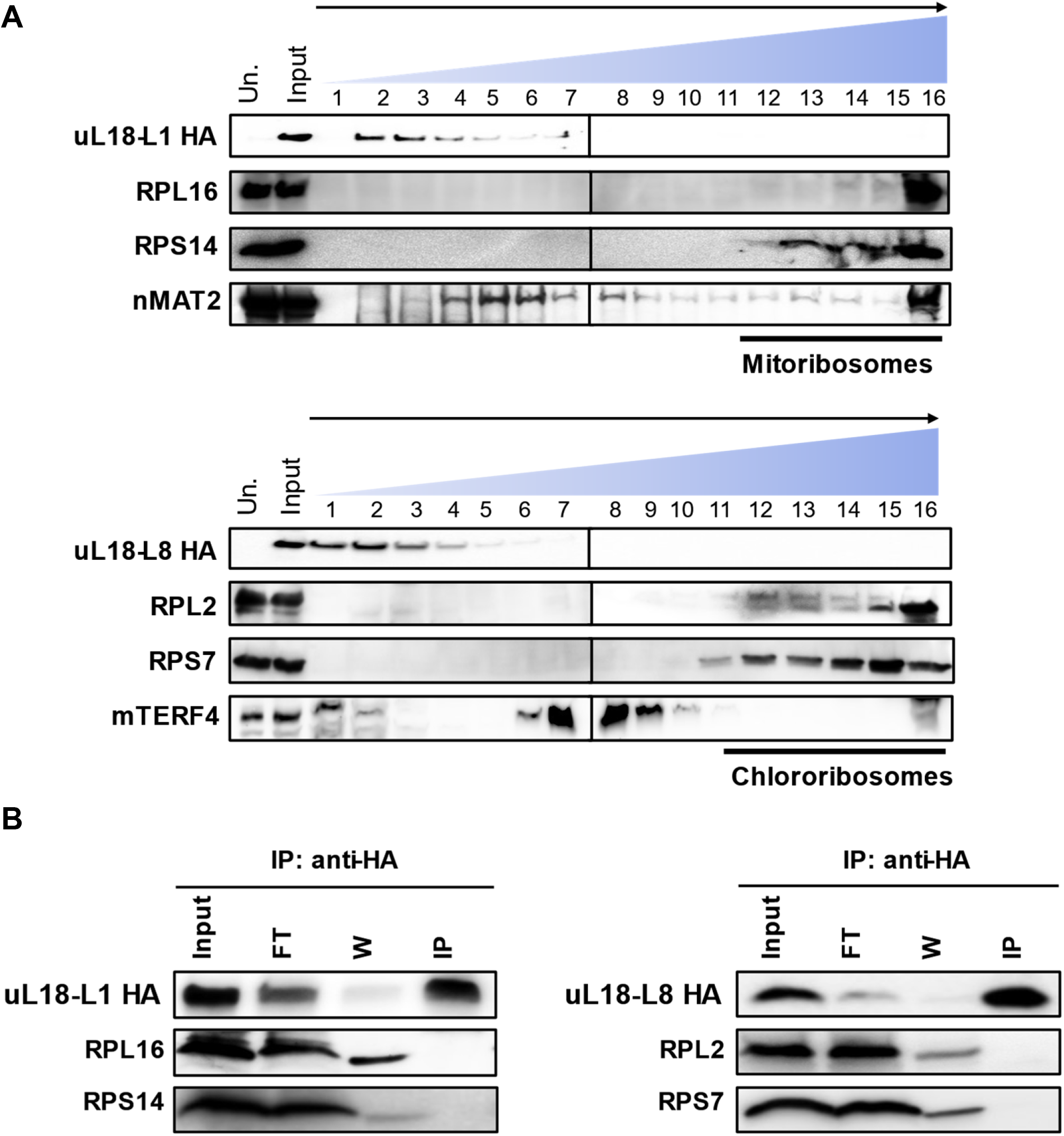
uL18-L1 and uL18-L8 do not take part in ribosomes. (A) Arabidopsis cell extracts prepared from cell lines over-expressing the indicated *uL18-L* genes were loaded on 10-30% sucrose gradients. Immunoblots of recovered fractions were probed with antibodies as detailed in Figure 7. Un.: untransformed cells. (B) Immunoprecipitation of uL18-L1-3HA and uL18-L8-3HA. Total extracts from transgenic Arabidopsis cell lines expressing uL18-L1-3HA or uL18-L8-3HA were used for immunoprecipitation with anti-HA antibody. Equal volumes of input, flow-through (FT), wash (W) and immunoprecipitated (IP) fractions were analyzed by immunoblot analysis using the indicated antibodies.

**Supplemental Table S2:** List of antibodies used in this study.

| **Antibody** | **Host** | **Dilution** | **Source** |
| --- | --- | --- | --- |
| AOX1/2 | Rabbit | 1 : 1,000 | Agrisera (AS04 054) |
| ATPC | Rabbit | 1 : 5,000 | Jorg Meurer (Ludwig-Maximilians-Universitat, Germany) |
| ATPβ | Chicken | 1 : 5,000 | Agrisera (AS05 085) |
| CA2 | mouse | 1 : 5,000 | Mariano Perales *et al.*, 2005 |
| Cox1 | Mouse | 1 : 1,000 | Meyer E. *et al.*, 2018 |
| Cox2 | Rabbit | 1 : 1,000 | Agrisera (AS04 053A) |
| Cytochrome C | Rabbit | 1 : 5,000 | Agrisera (AS08 343A) |
| HA (clone 3F10) | Rat | 1 : 500 | Roche (11867423001) |
| Nad9 | Rabbit | 1 : 5,000 | Lamattina L. *et al.*, 1993 |
| nMAT2 | Rabbit | 1 : 200 | Keren I. *et al.*, 2009 |
| PetA | Rabbit | 1 : 2,000 | Agrisera (AS08 306) |
| Plastocyanin | Rabbit | 1 : 2,000 | Agrisera (AS06 141) |
| Porin | Mouse | 1 : 500 | David Day (Western Australia University) |
| PsaD | Rabbit | 1 : 1,000 | Agrisera (AS09 461) |
| PsbD | Rabbit | 1 : 5,000 | Agrisera (AS06 146) |
| RISP | Rabbit | 1 : 5,000 | Chris C. *et al.*, 2010 |
| RPL16 | Rabbit | 1 : 1000 | Agrisera (AS15 3069) |
| RPL2 | Rabbit | 1 : 3,000 | Agrisera (AS15 2876) |
| RPS1 | Rabbit | 1 : 1,000 | Agrisera (AS15 2875) |
| RPS12 | Rabbit | 1 : 10,000 | Agrisera (AS12 2114) |
| RPS14 | Rabbit | 1 : 1000 | Agrisera (AS16 4094 ) |
| RPS7 | Rabbit | 1 : 3000 | Agrisera (AS15 2877) |
| Tubulin | Mouse | 1 : 5,000 | Sigma (T5168 ) |
| ZM-mTERF4 | Rabbit | 1 : 3,000 | Hammani K. *et al.*, 2014 |

**Supplemental Table S4:** Pigment content and chlorophyll fluorescence parameters in wild-type and *ul18-l8* mutant plants. Measurements of chlorophyll fluorescence parameters were performed with the PSI Open FluorCam FC 800 on dark-adapted plants. For 1-qP, qL, NPQmax measurements, an actinic light treatment (420 μmol photons.m^−2^.s^−1^) was performed for 500s. After termination of actinic light, NPQrelax is determined after 280s of dark. Data are presented as means ± SD (n=4). Grouping information using the Fisher Method and 95% confidence is given by bold lowercase letters.

| Parameter | Col-0 |  | *ul18-l8-1* |  | *ul18-l8-2* |  |
| --- | --- | --- | --- | --- | --- | --- |
| Water content (% fresh weight) | 90.94 ± 0.29 | **a** | 92.52 ± 1.55 | **a** | 92.37 ± 0.37 | **a** |
| Total chlorophyll (µg/mg fresh weight) | 0.632 ± 0.081 | **a** | 0.540 ± 0.077 | **a** | 0.541 ± 0.096 | **a** |
| Carotenoids (µg/mg fresh weight) | 0.086 ± 0.015 | **a** | 0.075 ± 0.009 | **a** | 0.076 ± 0.012 | **a** |
| Chlorophyll a/b | 2.37 ± 0.07 | **a** | 2.28 ± 0.05 | **a** | 2.248 ± 0.22 | **a** |
| Fv/Fm | 0.837 ± 0.012 | **a** | 0.743 ± 0.064 | **b** | 0.739 ± 0.022 | **b** |
| qL | 0.755 ± 0.073 | **a** | 0.451 ± 0.777 | **b** | 0.476 ± 0.009 | **b** |
| NPQ max | 2.127 ± 0.287 | **a** | 2.622 ± 0.257 | **b** | 2.589 ± 0.186 | **b** |
| NPQ relax | 0.666 ± 0.130 | **a** | 0.690 ± 0.211 | **a** | 0.673 ± 0.108 | **a** |

